# Artifact-free, high-temporal-resolution *in vivo* opto-electrophysiology with microLED optoelectrodes

**DOI:** 10.1101/622670

**Authors:** Kanghwan Kim, Mihály Vöröslakos, John P. Seymour, Kensall D. Wise, György Buzsáki, Euisik Yoon

## Abstract

The combination of *in vivo* extracellular recording and genetic-engineering-assisted optical stimulation is a powerful tool for the study of neuronal circuits. Precise analysis of complex neural circuits requires high-density integration of multiple cellular-size light sources and recording electrodes. However, high-density integration inevitably introduces stimulation artifact. We present minimal-stimulation-artifact (miniSTAR) µLED optoelectrodes that enable effective elimination of stimulation artifact. A multi-metal-layer structure with a shielding layer effectively suppresses capacitive coupling of stimulation signals. A heavily-boron-doped silicon substrate silences the photovoltaic effect induced from LED illumination. With transient stimulation pulse shaping, we reduced stimulation artifact on miniSTAR µLED optoelectrodes to below 50 µV_pp_, much smaller than a typical spike detection threshold, at optical stimulation of > 50 mW mm^-2^ irradiance. We demonstrated high-temporal resolution (< 1 ms) opto-electrophysiology without any artifact-induced signal quality degradation during *in vivo* experiments. MiniSTAR µLED optoelectrodes will facilitate functional mapping of local circuits and discoveries in the brain.

## Introduction

A brain is made up of densely populated neurons. Analysis of neuronal communication requires simultaneous high-resolution recording and neuron-specific perturbation of circuit components under controlled conditions. The combination of genetic engineering-assisted optical stimulation and massively parallel electrical recording of neuronal activities (opto-electrophysiology) is a promising tool for studying neuronal circuits in behaving animals^1^. A number of devices^2-11^ have been introduced for the past few years for *in vivo* opto-electrophysiology. For high-resolution *in vivo* opto-electrophysiology, a micromachined silicon multi-electrode-array structure also known as the Michigan Probe^12,13^ has been widely utilized^6-11^. These silicon optoelectrodes take advantage of the planar profile of the Michigan Probe platform and accommodate multiple light sources in the vicinity of high-density recording electrode arrays. This compact configuration provides the capability to electrically record activity of sets of neurons at high spatial resolution while optically stimulating a portion of the recorded neurons.

An undesirable feature of many of these devices is stimulation artifact. With its magnitude often an order of magnitude larger than those of underlying neuronal signals, the stimulation artifact may mask neuronal signals and prevent temporally precise recording of neuronal responses^14,15^. In order to enable precise detection of neuronal activities, the magnitude of the stimulation artifact should be reduced to lower than a threshold voltage level for neuronal activity detection. Typically, the threshold level is set as a few integer multiples (often 5 ×) of the root-mean-square value of background noise^16,17^. To keep the artifact magnitude lower than the threshold level, optical stimulation had been limited to slowly-changing, low-frequency pulses, such as slow (< 10 Hz) sine waves^11^ or trapezoidal pulses with a long (> 10 ms) rise time^18^. These slowly-changing optical stimulation protocols, however, are not suitable for many neuroscience experiments in which high-speed neuromodulation is required, such as those in closed-loop experimental setups^19^. An ideal optoelectrode should therefore provide optical stimulation with temporal resolution higher than the duration of the neuronal activities while keeping the stimulation artifact magnitude lower than a spike detection threshold.

We present minimal-stimulation-artifact (miniSTAR) µLED optoelectrode and report the engineering schemes that enabled artifact-free optical stimulation. We extensively characterized various forms of optical-stimulation-induced artifacts, including photoelectrochemical effects (PEC)^20-25^, electromagnetic interference (EMI)^6,26,27^, and photovoltaic effects (PV)^28,29^. MiniSTAR optoelectrodes utilize monolithically integrated neuron-sized LEDs^11^ for high spatial resolution optical stimulation of target neurons. The multi-metal-layer structure on the miniSTAR optoelectrode suppresses EMI-induced stimulation artifact, and the heavily-boron-doped silicon substrate eliminates PV-induced artifact. Additionally, transient pulse shaping control reduces the magnitude of residual stimulation artifact on all recording channels to < 50 µV peak-to-peak (µVpp) without compromising the temporal resolution of optical stimulation. With an *in vivo* experiment using a miniSTAR optoelectrode implanted in a mouse brain, we demonstrate the absence of distortion in the recorded neuronal signals during precise *in situ* optical stimulation.

## Results

### Fabrication and characterization of miniSTAR µLED optoelectrodes

We fabricated miniSTAR µLED optoelectrodes (Fig. 1a) using microfabrication techniques adapted from those used for the fabrication of the family of Michigan optoelectrodes including one-metal-layer µLED optoelectrodes^11^. Figure 1b describes the simplified device fabrication flow. MiniSTAR optoelectrodes were fabricated using gallium-nitride-on-silicon (GaN-on-Si), gallium nitride/indium gallium nitride multi-quantum-well (GaN/InGaN MQW) LED wafers with heavily boron-doped silicon (p^+^- Si, N^A^ ≈ 1 × 10^20^ cm^-3^) substrates. In order to reduce EMI-induced stimulation artifact, metal traces for LED drive signals (LED interconnects) and those for recorded neuronal signals (recording electrode interconnects) were placed in two different metal layers separated from each other by a ground-connected shielding layer (Fig. 1c, top), forming a multi-metal-layer structure. A heavily-boron-doped substrate was chosen to suppress diffusion of optically generated electron-hole pairs and, as a result, to reduce PV-induced stimulation artifact (Fig. 1c, bottom). First, LED mesa structures were formed on GaN/InGaN MQW layer, and the LED interconnects were defined on the first metal layer. After passivating the surface of the LEDs, the EMI shielding layer was defined on the second metal layer and the recording electrode interconnects were defined on the third metal layer. Neural signal recording electrodes were then formed by depositing electrode material (iridium) on top. Finally, the entire wafer was thinned down to 30 µm and the miniSTAR optoelectrodes were released from the silicon wafer. Released miniSTAR optoelectrodes were assembled on printed circuit boards (PCBs) that provide connections to a neuronal signal recording IC and an LED driver (Fig. 1d). Figure 1e shows a microphotograph of a tip of the fabricated miniSTAR optoelectrode. The dimensions of exposed surface area of each µLED and recording electrodes are 10 × 15 µm and 11 × 13 µm (W × L), respectively.

**Figure 1:**
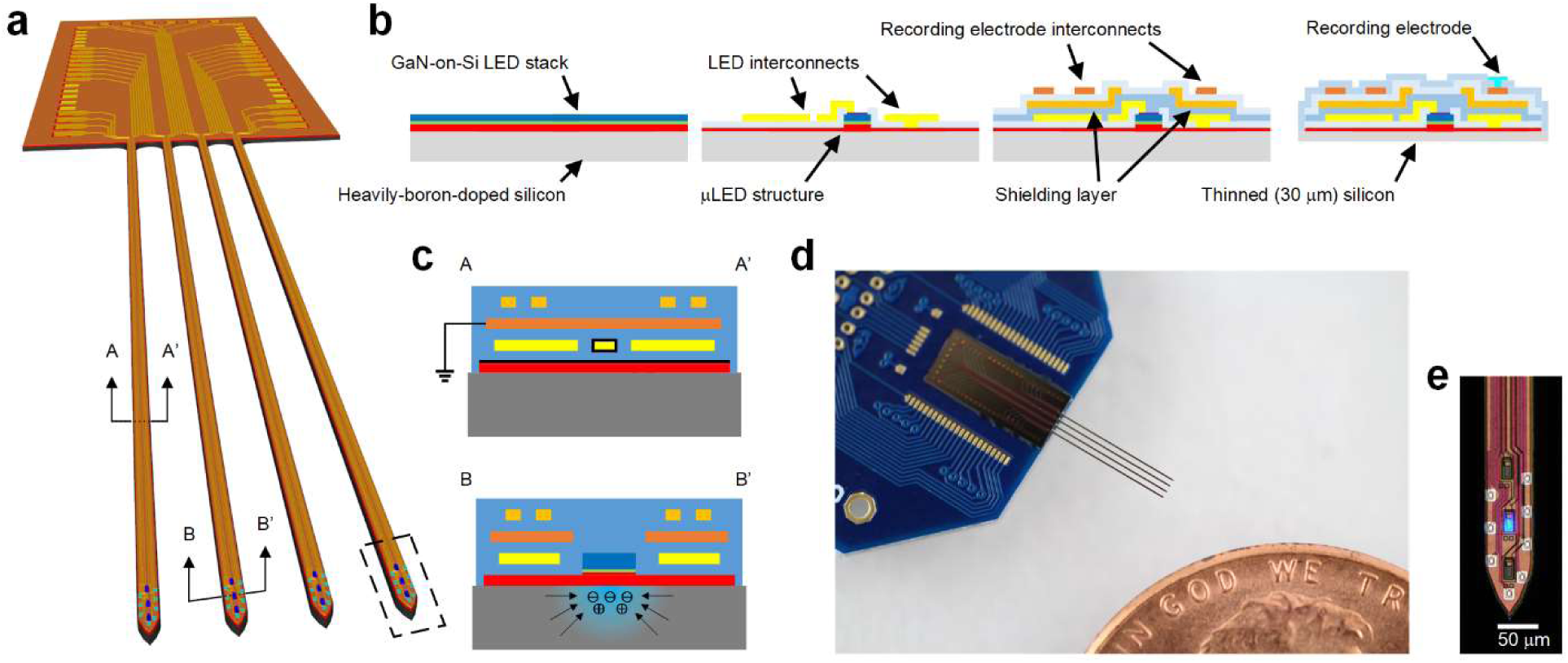
MiniSTAR optoelectrode. (**a**) Schematic illustration of a miniSTAR µLED optoelectrode. (**b**) Simplified miniSTAR optoelectrode fabrication process. An illustrative cross-section containing only one LED and one recording electrode is shown. (**c**) Cross-sectional schematic diagrams of a shank of miniSTAR optoelectrode, showing sources of stimulation artifact (EMI and PV effect for top and bottom, respectively) and methods for reduction of stimulation artifact. (**d**) Photograph of a miniSTAR optoelectrode mounted on a PCB. (**e**) Microphotograph of a tip of a miniSTAR optoelectrode, on which eight recording electrodes, three LEDs, LED interconnects, shielding layer, and recording electrode interconnects are shown.

After fabricating the miniSTAR optoelectrodes, we characterized performance of the LEDs and recording electrodes and confirmed that they are suitable for *in vivo* opto-electrophysiology. Optical power equivalent to greater than 1 mW mm^-2^ of irradiance is considered a threshold for activation of channelrhodopsin-2 (ChR2)^9,11^. LEDs on miniSTAR optoelectrodes generated a radiant flux of 150 nW, equivalent to an irradiance of 1 mW mm^-2^ at the surface when voltage of 2.86 ± 0.02 V (mean ± SD, n = 22) was applied across their terminals. The LEDs were capable of generating 50 mW mm^-2^ at the surface (7.5-µW radiant flux) at 3.46 ± 0.10 V, which is more than sufficient for activation of ChR2-expressing cells further away from the LED surface. We confirmed that the effect of substrate doping density on the electrical and optical characteristics of the fabricated LEDs is not as significant as the die-to-die variation in a wafer (Supplementary Figure 1). The impedance magnitude and phase of the recording electrodes were measured as 1.15 ± 0.07 MΩ and −68.33 ± 5.11 ° at 1 kHz (n = 54, mean ± SD), respectively, acceptable for high-quality *in vivo* extracellular recordings^30^.

### Reduction of EMI-induced artifact

EMI is inevitable in a system where a source of high-voltage, fast-changing signal is located in close proximity to a signal-carrying trace connected to a high-impedance load. Previous µLED optoelectrodes^30^ contained only one metal layer on which all the interconnects that carry optical stimulation signals as well as those carrying recorded neural signals were densely integrated. Therefore, mutual capacitances between the traces of two signal types were high, and, in turn, the recording interconnects were highly susceptible to EMI from LED drive signals. Moreover, the n-GaN layer that forms the common cathode of all the µLEDs on the optoelectrode was directly underneath the interconnects and acted as another significant source of EMI. FEM simulations of electrostatic potential distribution in the one-metal-layer µLED optoelectrode (Supplementary Results and Supplementary Figure 2c) showed significant voltage coupling from LED interconnects (−48.96 dB) as well as from the n-GaN layer (−0.06 dB).

We observed significant suppression of EMI-induced stimulation artifact with the integration of a shielding layer. We implemented the triple-metal-layer structure on µLED optoelectrode and dedicated a layer between the stimulation and recording interconnects as a shielding layer (Supplementary Figure 2d). Triple-metal-layer (shielded) µLED optoelectrodes were fabricated on the same GaN-on-Si LED wafer on which one-metal-layer µLED optoelectrodes were fabricated, which had a lightly boron-doped silicon substrate (N^A^ ≈ 5 × 10^16^ cm^-3^). We compared stimulation artifacts between one-metal-layer µLED optoelectrodes and shielded µLED optoelectrodes while turning on and off µLEDs *in vitro*.

Figure 2 shows the magnitude of the transient artifact (peak-to-peak) and the wideband and highpass filtered waveforms of artifacts resulting from optical stimulation. One-metal-layer µLED optoelectrodes showed a high transient magnitude (> 1 mVpp) in most recording sites regardless of the amount of optical power generated from the LEDs (Fig. 2b). The shape of wideband stimulation artifacts (Fig. 2c, left) suggests strong EMI-induced artifact. The weak dependence of stimulation artifact on optical power suggests that voltage coupling from the n-GaN substrate, whose voltage does not greatly change as a function of the LED signal voltage, greatly contributes to the EMI-induced stimulation artifact. On the other hand, shielded optoelectrodes showed significantly lower artifact (100 - 400 µVpp) which gradually increases at larger irradiance (Fig. 2e). It was also verified from both the wideband (Fig. 2f, left) and the highpass filtered waveforms (Fig. 2f, right) that the EMI-induced artifact has been greatly reduced. This result was consistent with expectation from the FEM simulations (Supplementary Figure 2c). Significant reduction of stimulation artifact on shielded optoelectrodes was observed at all irradiance under test (Fig. 2g). At 75-mW mm^-2^ irradiance (radiant flux of 11.5 µW), we achieved 5.2-fold reduction in stimulation artifact (from 2477.75 ± 1733.83 to 474.59 ± 146.26 µVpp, n = 75 and 67, respectively).

**Figure 2:**
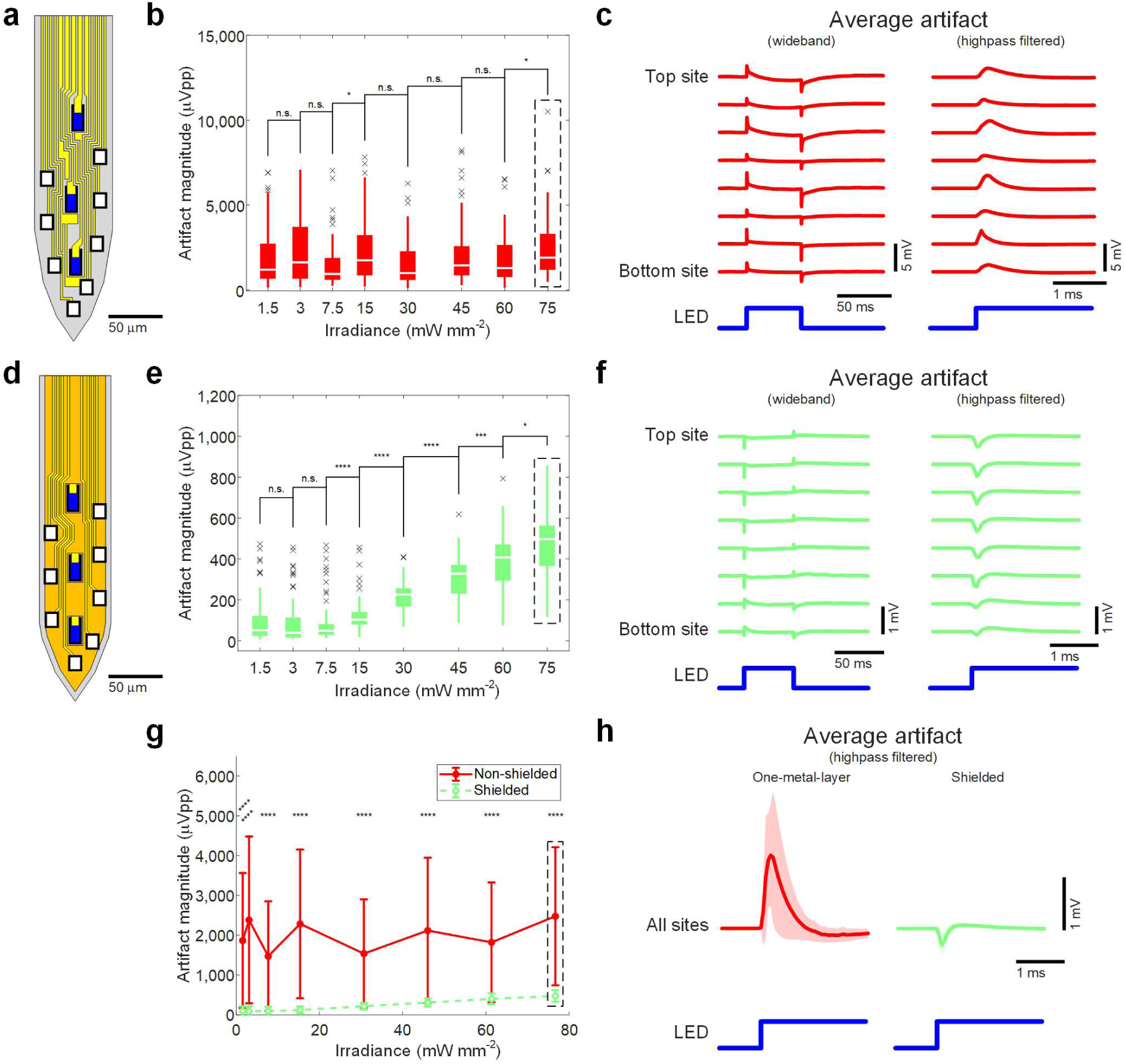
Reduced EMI-induced artifact. (**a**) Schematic illustration of the tip of one-metal-layer (non-shielded) µLED optoelectrode. Blue rectangles indicate LEDs, white rectangles the recording electrodes, and yellow polygons interconnects. (**b**) Peak-to-peak magnitude of highpass filtered stimulation artifact recorded on non-shielded µLED optoelectrodes. Data from channels corresponding to all electrodes on the shank on which a LED was turned on are plotted. Boxes indicate interquartile ranges, white lines medians, whiskers non-outlier extreme values, and black x marks outliers. (**c**) Mean waveforms of stimulation artifact recorded on non-shielded µLED optoelectrodes, from channels that correspond to electrodes on different locations on the tips. LED drive signal with resulting LED surface irradiance of 75 mW mm^-2^ was used. (**d**) Schematic illustration of the tip of shielded µLED optoelectrodes. The color scheme is identical to that of part *a*, except for an additional color, gold, to indicate shielding layer. (**e**) Peak-to-peak magnitude of highpass filtered stimulation artifact recorded on shielded µLED optoelectrodes. (**f**) Mean waveforms of stimulation artifact recorded on shielded µLED optoelectrodes. LED drive signal with resulting LED surface irradiance of 75 mW mm^-2^ was used. (**g**) Comparison of mean peak-to-peak magnitudes of highpass filtered stimulation artifact recorded on the shielded (green) and non-shielded (red) µLED optoelectrodes. Error bars indicate one standard deviation. (**h**) Mean highpass filtered waveforms whose mean peak-to-peak magnitudes are shown in part **g**, inside the rectangle with black dashed lines. Shaded regions show one standard deviation away from the mean. Mean (± SD) peak-to-peak magnitudes are 2477.8 (± 1733.83) for non-shielded µLED optoelectrodes (n = 75) and 474.6 (± 146.26) for shielded µLED optoelectrodes (n = 67). Results of statistical tests are provided in Supplementary Table 2.

### Elimination of PV-induced artifact

Although EMI-induced artifact was greatly suppressed with introduction of the shielding layer, the magnitude of the residual artifact was still high and should be further eliminated below that of typical neuronal spikes (∼ 100 µVpp). Interestingly, we noticed that the polarity of stimulation artifact at the rising and falling edges of LED drive pulses were inverted on the shielded µLED optoelectrodes. As can be seen in Fig. 2c, the transient artifact on one-metal-layer µLED optoelectrodes has the same positive polarity as that of the LED drive signal, forming an inverted-‘v’ (or ‘^’) shaped waveform. However, on the shileded µLED optoelectrodes (Fig. 2f and h), the polarity of the transient artifact was inverted, making a v-shaped waveform. Inversion of the polarity of the transient artifact suggested that the residual artifact could result from a different source other than EMI.

We hypothesized that the source of v-shaped stimulation artifacts is photovoltaic (PV) effects in the silicon substrate and confirmed our hypothesis with a few experiments. First, we observed the waveform of signals recorded on electrodes while exposing the µLED optoelectrodes to external optical illumination. Using a focused beam at a wavelength similar to that of the light generated from µLEDs (λ_peak_ ≅ 470 nm), we illuminated tips of the shielded µLED optoelectrodes. The shape of induced voltage signals was identical to that of the stimulation artifact observed on the optoelectrodes (Supplementary Figure 3). The identical shape suggested that the artifact is truly optically induced, not resulting from EMI. We repeated the experiment using electrode arrays fabricated on non-silicon substrates: GaN-on-sapphire wafer and soda lime glass. We did not observed any v-shaped stimulation artifacts on electrodes on both substrates (Supplementary Figure 4), verifying that the artifact is due to neither photoelectrochemical (PEC) effects on the electrodes nor PV-induced artifact on the GaN layer. With the exclusion of PEC effects and PV effect from the GaN layer, the only remaining source of potential light-induced artifact is the PV effects from the silicon substrate.

A few experimental studies in the past reported that light-induced noise on silicon electrode arrays can be reduced with use of heavily-doped substrate^31,32^. Heavy doping of semiconductor greatly reduces carrier lifetimes^33,34^ and diffusion lengths of free carriers, which supposedly contributes to the amount of dipole-induced voltage^31^. Therefore, PV-induced stimulation artifacts should be suppressed with heavy doping of the silicon substrate. We conducted FEM simulations of optically induced voltage generation in silicon substrates and verified that the voltage is reduced with heavy substrate doping. We built a model of the silicon substrate and calculated the optically-induced voltage generation while changing doping concentrations (Supplementary Methods). The results suggest that highly a boron-doped silicon substrate can greatly reduce the magnitude of optically induced voltage and as a result suppress PV-induced artifact (Supplementary Results and Supplementary Figure 5).

In order to verify the effect of doping density on the magnitude of PV-induced stimulation artifact, we fabricated three groups of shielded µLED optoelectrodes using GaN-on-Si GaN/InGaN MQW LED wafers with different silicon substrates: float-zone grown silicon substrate (FZ-Si, N^A^ ≈ 4 × 10^12^ cm^-3^), lightly boron-doped silicon substrate (p^-^-Si, N^A^ ≈ 5 × 10^16^ cm^-3^), and heavily boron-doped silicon substrate (p^+^-Si, N^A^ ≈ 1 × 10^20^ cm^-3^). Fig. 3a shows that the magnitude of stimulation artifact measured on the optoelectrodes fabricated using wafers with FZ-Si and p^-^-Si substrates increases as a function of irradiance (FZ-Si: 109.59 ± 80.61 µVpp at 1.5 mW mm^-2^ increasing to 569.33 ± 129.00 µVpp at 75 mW mm^-2^, p^-^-Si: 99.25 ± 116.01 µVpp at 1.5 mW mm^-2^ increasing to 474.59 ± 146.26 µVpp to at 75 mW mm^-2^, mean ± SD). On the other hand, the stimulation artifact magnitude on devices with p^+^-Si substrate did not show significant change (133.04 ± 121.99 µVpp at 1.5 mW mm^-2^ to 146.05 ± 143.4 µVpp at 75 mW mm^-2^, mean ± SD). The magnitude of stimulation artifact as a function of irradiance and substrate doping density (Fig. 3b) was similar to that expected from FEM simulation (Supplementary Figure 5d). Figure 3c shows the waveforms of stimulation artifact measured on the optoelectrodes of each group. It can be seen that even with high-intensity illumination (11.5 µW, 75 mW mm^-2^), the mean magnitude of stimulation artifact was below 200 µVpp, suggesting that PV-induced stimulation artifact was effectively reduced by use of heavily-boron-doped silicon substrate.

**Figure 3:**
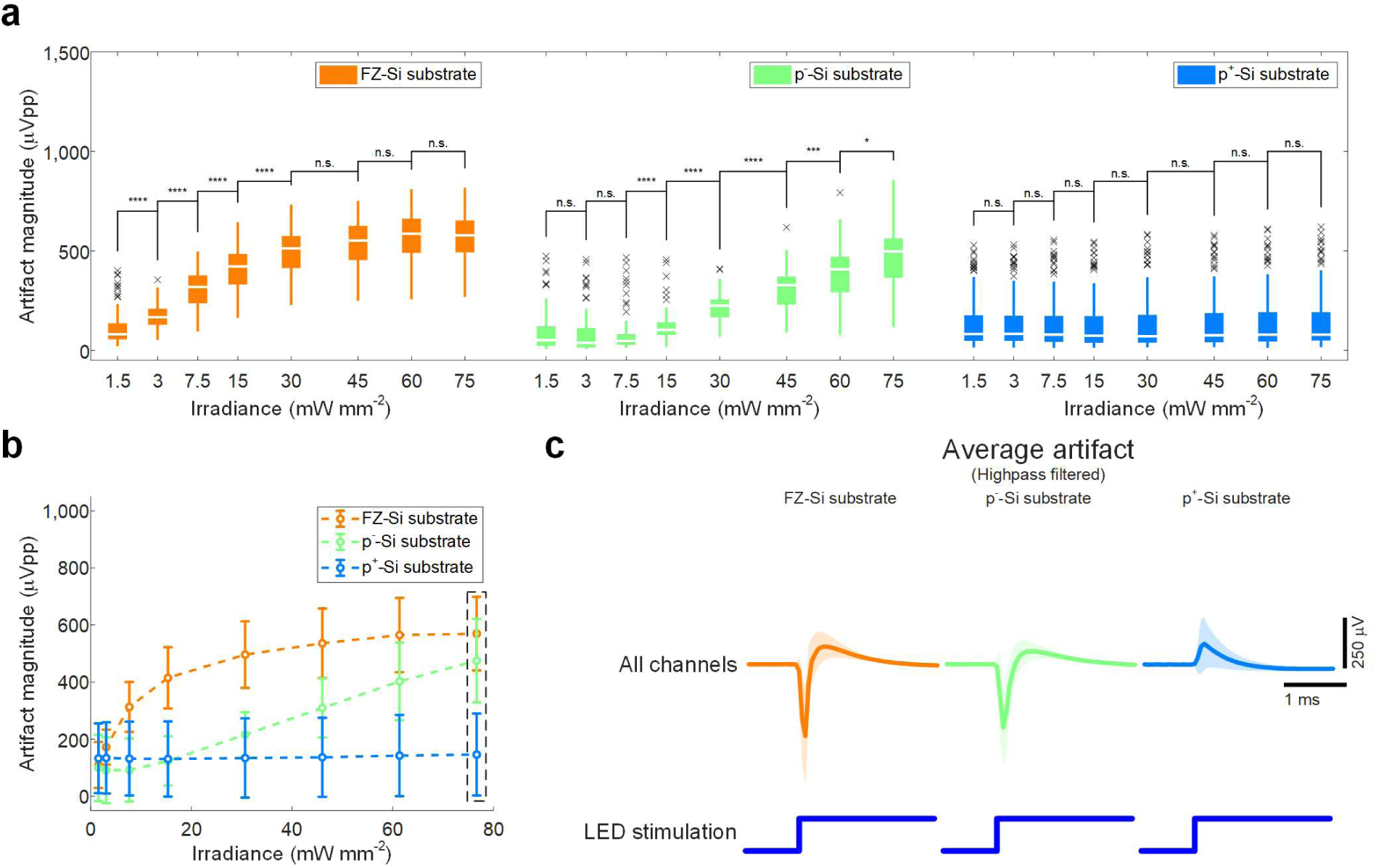
Eliminated PV-induced artifact. (**a**) Peak-to-peak magnitude of highpass filtered stimulation artifact recorded on shielded µLED optoelectrodes with different substrate doping densities. Data from channels corresponding to all electrodes on the shank on which an LED was turned on are plotted. Boxes indicate interquartile ranges, white lines medians, whiskers non-outlier extreme values, and black x marks outliers. (**b**) Comparison of mean peak-to-peak magnitude of highpass filtered stimulation artifact whose distribution is shown in part **a**. Error bars indicate one standard deviation. (**c**) Mean highpass filtered waveforms whose mean peak-to-peak magnitudes are shown in part **b**, inside the rectangle with black dashed lines. Shaded regions show one standard deviation away from mean. The mean (± SD) peak-to-peak magnitudes are 569.33 (± 129.00), 474.59 (± 146.26), and 146.05 (± 143.40) µVpp for devices with FZ-Si substrate (n = 124), p^-^-Si substrate (n = 67), and p^+^-Si substrate (n = 151), respectively. Results of statistical tests are provided in Supplementary Table 2.

We confirmed elimination of the PV-induced stimulation artifact by inspecting the shape and the magnitude of stimulation artifact waveforms recorded from electrodes at different locations on µLED optoelectrodes (Fig. 4). Figure 4b shows the magnitude of stimulation artifact recorded from channels that correspond to the electrodes marked in Fig. 4a. The artifact waveform recorded from each channel is presented in Fig. 4c. It is interesting to note that, while the v-shaped waveform in stimulation artifact was observed in the recordings from optoelectrodes with FZ-Si and p^-^-Si substrates, we no longer observed the v-shape in the optoelectrodes with p^+^-Si substrates. The absence of the characteristic v-shaped waveform confirms that PV-induced stimulation artifact has been eliminated on the optoelectrodes with p^+^ silicon substrate.

**Figure 4:**
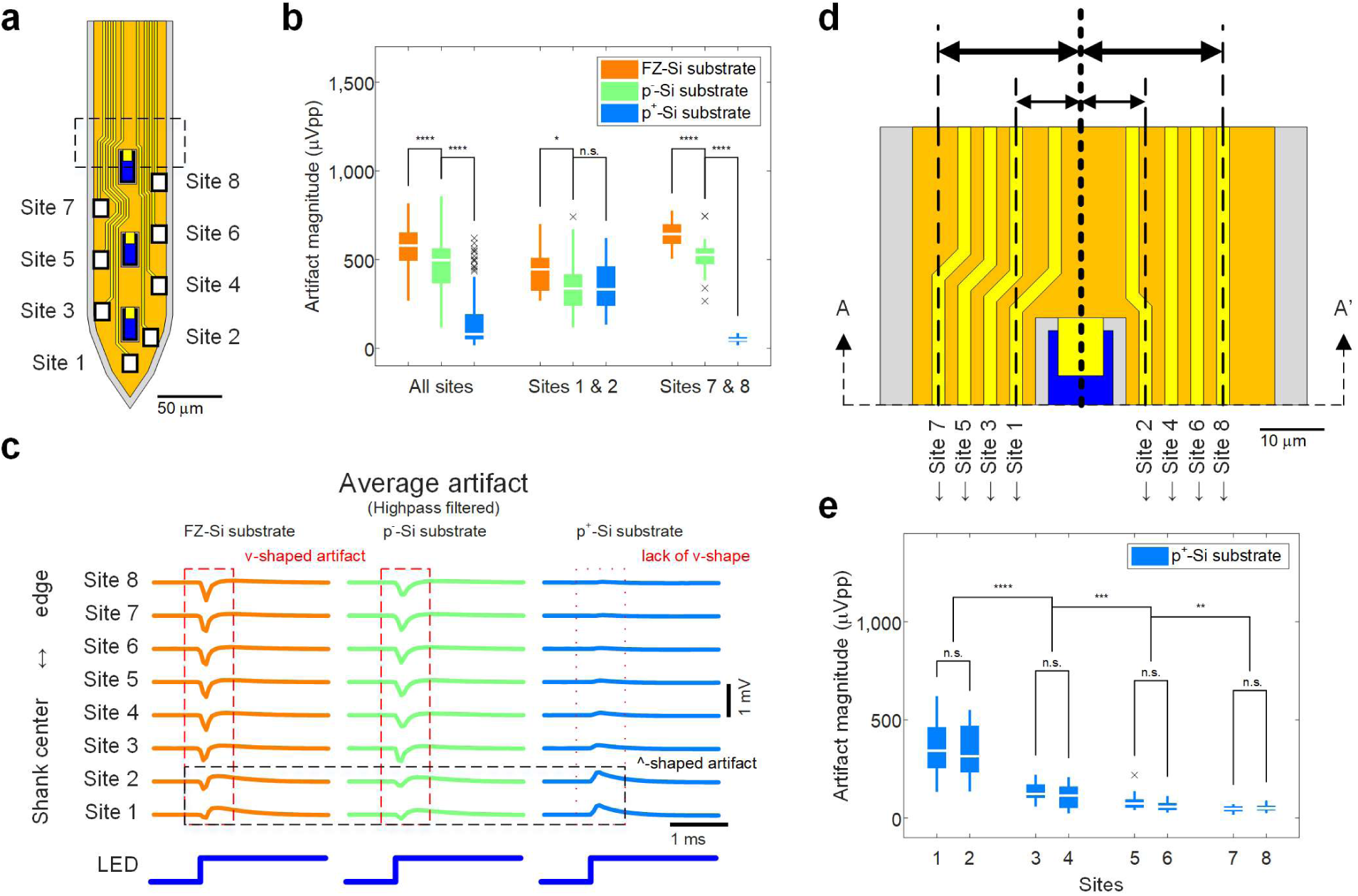
Location dependence of residual artifact. (**a**) Schematic illustration of the tip of shielded µLED optoelectrode. (**b**) Peak-to-peak magnitude of highpass filtered stimulation artifact recorded on shielded µLED optoelectrodes with different substrate doping densities. Data from channels corresponding to electrodes on the shank on which an LED was turned on are plotted. LED drive signal with resulting LED surface irradiance of 75 mW mm^-2^ was used. Boxes indicate interquartile ranges, white lines medians, whiskers non-outlier extreme values, and black x marks outliers. (**c**) Mean waveforms of stimulation artifact recorded on the shielded µLED optoelectrodes with different substrate doping densities, from channels that correspond to electrodes on different locations on the tips. LED drive signal with resulting LED surface irradiance of 75 mW mm^-2^ was used. (**d**) Magnified view of the region inside the rectangle with the black dashed lines on part **a**. The distances between the center of the interconnects and the center of an LED are shown. (**e**) Peak-to-peak magnitudes of highpass filtered stimulation artifact recorded from different channels on shielded µLED optoelectrodes with heavily boron-doped silicon substrate (miniSTAR optoelectrodes). LED drive signal with resulting LED surface irradiance of 75 mW mm^-2^ was used. Results of statistical tests are provided in Supplementary Table 2.

### Suppression of residual EMI-induced artifact by transient stimulation pulse shaping

Considering great reduction of both EMI- and PV-induced stimulation artifact, we refer to the shielded µLED optoelectrodes fabricated using LED wafer with p^+^ silicon substrate as minimal-stimulation-artifact (miniSTAR) µLED optoelectrodes. We quantified the amount of reduction in stimulation artifact from the implementation of shielding layers and the replacement of substrate with highly-boron doped silicon in miniSTAR optoelectrodes (Supplementary Figure 6). The magnitude of artifact was reduced by a factor of 5.2 in average only from the shielding layers (from 2477.75 ± 1733.83 to 474.59 ± 146.26 µVpp, at 11.5 µW, mean ± SD), and by a factor of 17 in average from both shielding and substrate replacement combined (to 146.05 ± 143.40 µVpp, at 11.5 µW, mean ± SD). However, the magnitude of stimulation artifact in a couple of recording sites (sites 1 and 2) was still high, as large as 200 - 300 µVpp, while those on some other sites (sites 7 and 8) were less than 50 µVpp (Fig 4e).

Location dependence of residual stimulation artifact revealed that the residual artifact is due to EMI resulting from imperfection in the shielding layer. The shieling layer on miniSTAR optoelectrode contains openings (or optical windows) on top of µLEDs for illumination. However, the optical windows allow the electric field generated from the LEDs to exit the shielding layer and makes the interconnects susceptible to EMI. Once the PV-induced artifact was removed in miniSTAR optoelectrodes, we observed emergence of ^-shaped waveforms (Fig. 4c), which is especially pronounced on sites 1 and 2. The magnitude of ^-shaped waveform is inversely proportional to the distance between the interconnect for each site and the optical window on the shielding layer (Fig. 4c-e). The polarity and the distance dependence of stimulation artifact waveforms suggest that this residual artifact is due to EMI originating from the LEDs that are exposed through optical windows on the shielding layer.

Additional suppression of residual artifact was achieved by transient pulse shaping of LED drive signal. We modified the slew rate of voltage pulses by changing the rise and fall times of the pulses. With sufficiently long rise time (t_rise_ > (2πF_s_)^-1^), the magnitude of higher-order harmonics of the coupled signal that contributes to the artifact ((πt_rise_)^-1^ < f < F_s_/2) is reduced by additional −20 dB/decade (Supplementary Figure 7). Figure 5 shows the peak-to-peak magnitude and waveforms of stimulation artifact recorded from the channels corresponding to the bottom two electrodes (sites 1 and 2) on the tip of miniSTAR optoelectrodes, which show the worst residual EMI-induced artifact. We observed significant reduction in stimulation artifact as we increased the rise time longer than 100 µs. At 50 mW mm^-2^ irradiance, the artifact magnitude was reduced below 200 µVpp (173.99 ± 55.76 µVpp, mean ± SD, for 1-ms rise time). In order to further reduce the slew rate of the voltage driving signal, we adjusted the low-level (or off-state) voltage in the stimulation pulse signals. We increased the low-level voltage to 2.8V, just below the lowest turn-on voltage of LEDs. The voltage required for irradiance of 50 mW mm^-2^ (radiant flux of 7.5 µW) is approximately 3.5 V. By adjusting the low-level voltage from 0 V to 2.8 V, we reduced the voltage swing from 3.5 V to 0.7 V and the slew rate by a factor of 5. We confirmed that the artifact magnitude can be reduced to 111.92 µVpp (SD = 55.76 µVpp) even without adjusting the rise time (Fig. 5b, V_low_ = 2.8 V, blue). With a 1-ms rise time and 2.8-V low-level voltage, the mean artifact magnitude was reduced to 46.53 µVpp (SD = 11.33 µVpp). In typical *in vivo* extracellular measurements, 100 µVpp is used as a spike detection threshold due to biological and environmental noise. Therefore, stimulation artifact with less than 50-µVpp magnitude can be considered nearly artifact-free.

**Figure 5:**
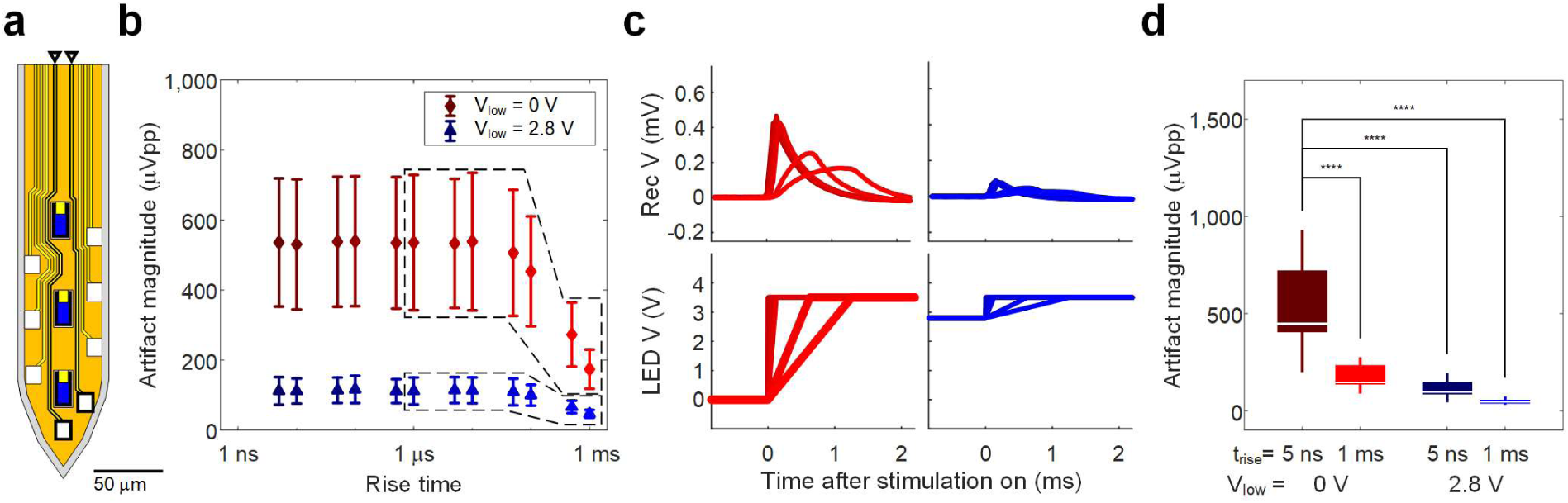
Effect of transient pulse shaping on residual artifact. (**a**) Schematic illustration of the tip of miniSTAR µLED optoelectrodes. Locations of the electrodes and interconnects from which the signals were recorded are indicated with rectangles with bold black lines and black arrowheads, respectively. (**b**) Mean peak-to-peak magnitude of highpass filtered stimulation artifact recorded from the channels indicated in part **a** for two different low-level voltages (V_low_ = 0 V and V_low_ = 2.8 V). High-level voltage of 3.5 V was used. X coordinates indicate the 10 - 90 % rise time of the pulse, and error bars indicate one standard deviation (n = 35). (**c**) Mean waveforms of recorded stimulation artifact, whose mean peak-to-peak magnitudes are shown inside the polygon with dashed lines in part **b**, and their input voltage signals. Stimulation artifact resulting from an input voltage signal is indicated with the same color. (**d**) Peak-to-peak magnitudes of highpass filtered stimulation artifact for a few selected conditions whose means are shown in part **b**. Boxes indicate interquartile ranges, white lines medians, and whiskers extreme values. Mean (± SD) peak-to-peak magnitudes are 535.80 (± 182.94), 173.99 (± 55.76), 111.92 (± 39.55), and 46.53 (± 11.33), from left to right. Results of statistical tests are provided in Supplementary Table 2.

### Validation of stimulation-artifact-free *in vivo* opto-electrophysiology

Following *in vitro* characterization, we demonstrated successful elimination of supra-threshold stimulation artifact *in vivo*. We implanted a miniSTAR µLED optoelectrode in the brain of a mouse and positioned its tips in the CA1 region of the hippocampus (Fig. 6a). Once spontaneous spikes and the characteristic high-frequency oscillations (ripples) were detected from the recording electrodes on a shank, each LED on the shank was turned on with varying powers to identify the optimal intensity of optical stimulation to alter the spiking activity of neurons (‘localized effect’) without inducing high-frequency oscillations due to synchronized firing of neuron populations^11^. Considering the typical duration of an action potential (< 2 ms), we used a rise/fall time of 1 ms to ensure maximum reduction of stimulation artifact without loss of the temporal resolution in optical stimulation.

**Figure 6:**
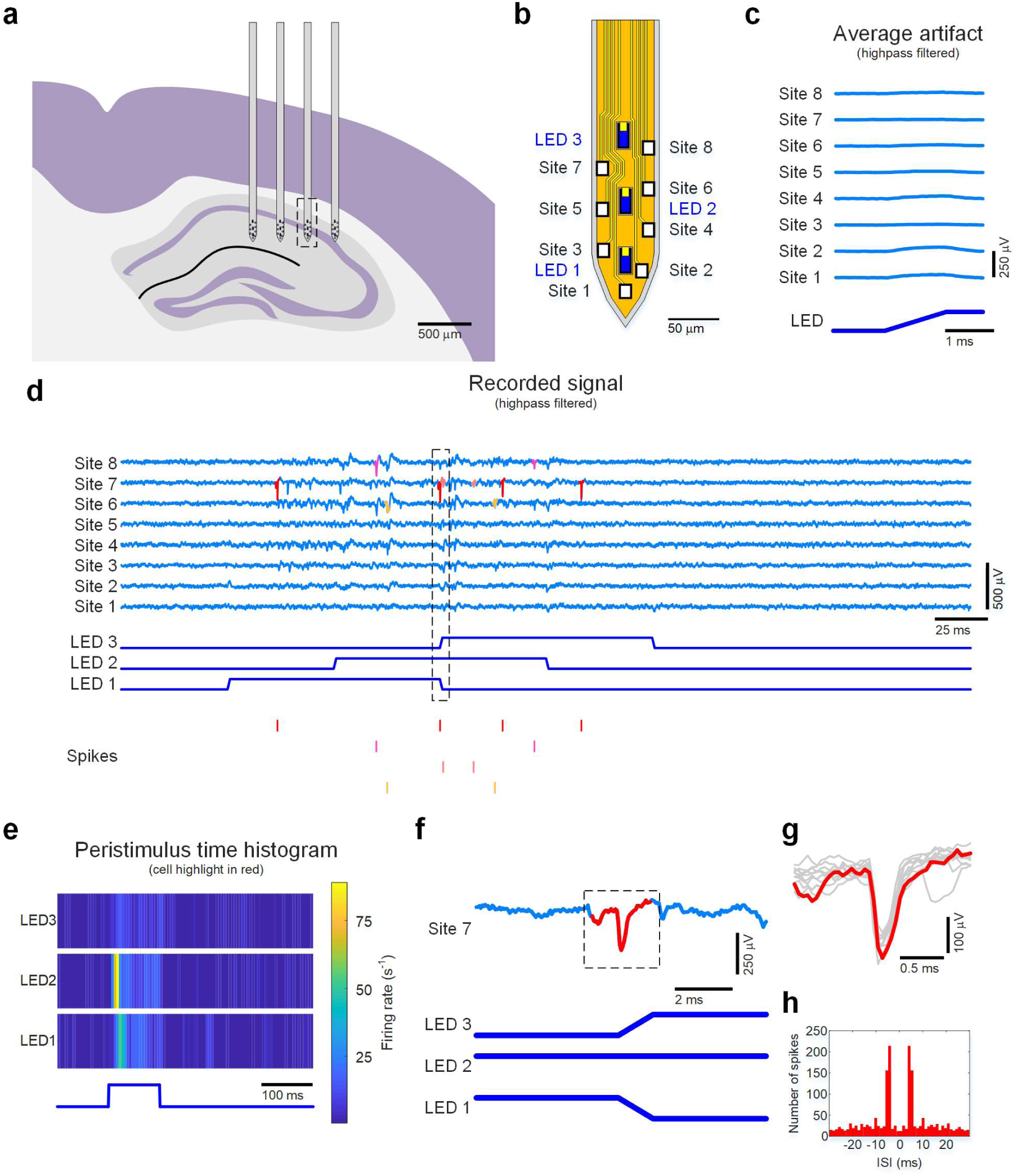
Stimulation-artifact-free *in vivo* opto-electrophysiology. (**a**) Schematic illustration of the location of implanted miniSTAR µLED optoelectrode inside the brain. The shank from which the data presented in parts c-h were collected is highlighted with a rectangle with dashed lines. (**b**) Schematic illustration of the tip of the miniSTAR optoelectrode. (**c**) Mean waveforms of stimulation artifact recorded on miniSTAR optoelectrodes, from channels that correspond to the electrodes on different locations on the tips. LED drive signal with resulting LED surface irradiance of 3 mW mm^-2^ at the surface of the LED (radiant flux of 460 nW) was used. (**d**) Traces of the recorded signals and the raster plots of some sorted spikes. Strong light-induced activities and lack of stimulation artifact on all the recording channels are shown. (**e**) Peristimulus time histograms of one of the identified cells whose spikes are highlight in red color in part **d**. (**f**) Magnified view of the signal recorded from Site 7 and the signals provided to the LEDs inside the region highlighted with a rectangle with black dashed line in part **d**. Note that the spike is not masked by either the offset of stimulation at site 1 or the onset of stimulation at site 3. (**g**) Magnified view of the spike inside the region highlighted with a rectangle with black dashed line in part **f**, overlaid on top of waveforms of ten spontaneous spikes from the same neuron. (h) Autocorrelogram of the spikes of the highlighted cell.

Stimulation with a 460-nW radiant flux (irradiance of 3 mW mm^-2^ at the surface of each µLED) induced strong light-induced responses in adjacent neurons. Optical stimulation with higher intensities induced high-magnitude (> 100 µVpp) population bursting of multiple cells^11^, preventing identification of single neurons and analysis of stimulation artifact. The mean waveform of signal the recorded from channels corresponding to electrodes at different locations on the shank (Fig. 6b) during the onset of the 3-mW mm^-2^ optical stimulation is shown in Fig. 6c. No supra-threshold (> 50 µVpp) stimulation artifact was observed from most channels.

After characterization of the stimulation artifact while driving individual LEDs, all three LEDs on the same shank were turned on and off in an interleaved manner (highlighted with a rectangle with black dashed lines in Fig. 6d). As shown in Fig. 6d, the series of optical pulses did not generate noticeable stimulation artifacts that would prevent either online detection of spikes or their offline spike sorting. With offline spike sorting^35^, we identified 6 putative pyramidal neurons with distinct spike waveforms in the vicinity of the shank on which the LEDs were activated. Further analysis of processed data identified a neuron which was clearly detected at the time stimulation offset of LED 1 and onset of LED 3 (Fig. 6f). No noticeable distortion of the spike waveform was present due to optical stimulation (Fig. 6g).

## Discussion

Results of this study the demonstrated the capability of high-spatiotemporal-resolution *in vivo* opto-electrophysiology with miniSTAR µLED optoelectrodes. We validated that the implementation multi-metal-layer structure, high-density boron doping of silicon substrate, and transient pulse shaping of stimulation waveform can effectively suppress stimulation artifact. A few non-ideal features in the fabricated miniSTAR optoelectrodes prevented the magnitude of the stimulation artifact from being further reduced. One imperfection is existence of optical windows on the shielding layer, which allow EMI generated from LEDs to reach their neighboring recording electrode interconnects. The other non-ideal factor is that the shielding layer has non-zero resistance. The shielding layer, especially near the tips of the shanks, is not strictly an ideal ground due to a resistive voltage drop through the thin-film metal layer it is made of. This resistive voltage drop would make the voltage of the shielding layer fluctuate as the voltage of LED interconnect changes, and the shielding layer itself could have acted as a source of EMI. These non-ideality resulted in less efficient reduction of EMI-induced artifact than that FEM electrostatic simulation predicted.

Residual EMI artifact might be able to be further suppressed with a few additional techniques. First, techniques to reduce the mutual capacitance between the recording electrode interconnect and the LED anode interconnect can be utilized. Increasing the distance between the recording electrode interconnects and the optical windows on the shielding layer (Fig. 4d) results in reduction of the mutual capacitance. Alternatively, a pair of ground-connected traces serving as shielding guards could have been placed between the optical windows and the recording electrode interconnects. However, these measures to reduce the mutual capacitance would inevitably increase the width of the shanks. Shank width is the main limiting factor for high-density scaling of the device which is required for larger-scale recording applications. Therefore, these options might not be considered optimal.

Methods to reduce electrode impedance might also be utilized for further reduction of EMI-induced artifact. The current carrying the capacitively coupled signal is divided between two branches in the signal recording circuit each of which is terminated with the amplifier load and the electrode (Supplementary Figure 8a). Therefore, lowering the electrode impedance would result in less current flowing through the amplifier load and thus reduction in the magnitude of the recorded voltage. In the data we collected using fabricated miniSTAR optoelectrodes, however, no obvious correlation between the electrode impedance and the magnitude of the stimulation artifact was observed (Supplementary Figure 8b). The small variance in the electrode impedance magnitude could have contributed to the lack of correlation. In future versions of miniSTAR optoelectrodes, electrode surface modification techniques such as site-level electrodeposition of conductive nanoparticles (e.g. Pt nanoparticle^36,37^ and PEDOT:PSS^38-40^) might be utilized to reduce the electrode impedance. The relationship between the electrode impedance and the stimulation artifact magnitude can then be studied in depth with groups of different miniSTAR optoelectrodes with significantly different electrode impedance ranges.

Transient stimulation pulse shaping complements the two engineering schemes of structural changes implemented in the miniSTAR optoelectrodes and effectively eliminates stimulation artifact. With nearly-zero amplitude artifact, miniSTAR optoelectrodes can be readily utilized for applications that require real-time event detection and closed-loop perturbation of neural circuits^1,19^. Still, the tradeoff between the temporal precision of optical stimuli and the amount of reduction in stimulation artifact should be taken into consideration. Most opsins have slow kinetics and will not provide sub-millisecond-precision responses regardless of the precision of optical stimulus. Short-pulse optical stimulation protocols, similar to electrical protocols utilizing a train of sub-millisecond pulses^41^, might be found useful in combination with some fast-responding opsins^42,43^. Generation of such short (< 2 ms) optical pulses may result in discernable signatures (> 50 µVpp) in the recordings. These potential supra-threshold-amplitude artifacts, however, can easily be subtracted^44^ from the recorded trace since they occur at predetermined times and display identical waveforms. Therefore, residual stimulation artifact would not significantly compromise the quality of recording.

In some applications, current-based LED driving might be more desirable than voltage-based driving. When driven with current pulses, the voltage changes across the two terminals of an LED would follow the I-V characteristics of the LEDs. Therefore, current driving allows setting the non-zero off-state voltage across an LED, typically just below the LED turn-on voltage (≈ 2.8 V). We validated that the effect of current-based driving of LEDs is similar to that with voltage-based driving of LEDs with non-zero low-level voltage (Supplementary Figure 9). We further tested pulses with three different rise- and fall-time shapes: trapezoidal, sinusoidal and sigmoidal. The shape of current pulses during on-to off-state transition did not significantly affect the magnitude of stimulation artifact. This result suggests that, if stimulation pulses have sufficiently low slew rate, smoothening of pulse edges does not necessary provide additional reduction in stimulation artifact magnitude.

Overall, our work demonstrates that stimulation artifact can be successfully suppressed using miniSTAR µLED optoelectrodes. This new device will allow performing high temporal resolution *in vivo* opto-electrophysiology for in-depth understanding of the interactions among the multiple components of neuronal circuits.

## Methods

### Shielded µLED optoelectrode fabrication and device assembly

Four-inch-diameter silicon wafers with different substrate boron doping densities (N_A_ ≈ 4 × 10^12^, 5 × 10^16^, and 1 × 10^20^ cm^-3^, respectively) with GaN/InGaN multi-quantum-well (MQW) LED layers epitaxially grown with metal-organic chemical vapor deposition (MOCVD) on top were purchased from Enkris Semiconductor (Suzhou, China). LED structures, including LED mesas, p- and n-GaN contacts and metallic interconnects, were first formed on the wafer. Repeated deposition of passivating dielectric layers and deposition of patterned metal layers formed additional metal layers. Consecutively, the top metal layer was passivated, and neural signal recording electrodes were defined. Finally, fabricated µLED optoelectrodes were thinned and released from the silicon wafer by double-sided plasma dicing process. The detailed fabrication process, including the list of tools used for each fabrication step, is provided in Supplementary Methods.

In order to prevent unwanted capacitive voltage coupling at assembly level, we used four-layer printed circuit boards (PCBs) on which the traces for recorded neuronal signals and LED drive signals are separated by two ground-connected internal layers. The optoelectrodes were mounted on the PCBs and were electrically connected to the PCBs by wirebonding contact pads on the backend of the optoelectrode to the gold pads on the PCBs. After wirebonding, exposed wires were potted with epoxy (EPO-TEK 353ND and 353NDT, Epoxy Technology, Billerica, MA), and connectors (Omnetics Connector Corp., Minneapolis, MN) as well as the ground and the reference wires were soldered to the PCBs to finalize assembly process.

### Characterization of the electrical and optical properties of the µLEDs

The electrical and optical properties of each µLED on the µLED optoelectrodes were characterized before *in vitro* and *in vivo* characterization of stimulation artifact. Both current-voltage (I vs.V) and the irradiance-voltage (E_e_ vs. V) characteristics were measured for each µLED. First, an optical measurement system consisting of an integrating sphere (FOIS-1, Ocean Optics, Largo, FL) and a spectrometer (Flame, Ocean Optics) was built. A sourcemeter (Keitheley 2400, Keithley Instruments, Cleveland, OH) was then connected across the anode and the cathode pins of an µLED on the connector. The tips of the optoelectrode were lowered until the whole shanks were completely inside the integrating sphere, ensuring that all the light generated from the µLED can be collected. The DC voltage across the LED anode and the cathode terminals were swept from 0 V to 4 V, and the resulting current and the spectral flux of the µLED were measured. The radiant flux was calculated by integrating the spectral flux over wavelengths from 350 nm to 600 nm, and the irradiance on the surface of the µLED was then calculated by dividing the radiant flux by the µLED’s surface area (150 µm^2^).

### Characterization of the electrical properties of the recording electrodes

The impedance (both the magnitude and the phase at 1 kHz) of each recording electrode on the µLED optoelectrode was measured using an Intan neural signal recording amplifier (RHD2132, Intan Technologies, Los Angeles, CA) in 1 × phosphate buffered saline (PBS) solution (prepared using 10 × PSB purchased from MP Biomedicals, Solon, OH). First, a small amount of PBS (approximately 100 mL of 1 × PBS solution) was poured into a small clear polystyrene container (530C-CRY, AMAC Plastic Products, Petaluma, CA). The µLED optoelectrode was lowered into the container until the bottom halves of the shanks (∼ 2.5 mm) were submerged in PBS. Exposed stainless steel tips at the loose ends of the ground and the reference wires were also submerged in PBS. A neuronal signal recording system (RHD2000, Intan Technologies) conducted electrode impedance measurements.

### Setup for *in vitro* characterization of LED-drive-induced artifacts

*In vitro* characterization was conducted in 1 × PBS solution in AMAC 530C-CRY container. An µLED optoelectrode was lowered into the container until the bottom halves of the shanks were submerged in PBS. The exposed stainless steel tips at the loose ends of the ground and the reference wires were also submerged in the PBS.

An Intan RHD2000 neuronal signal recording system, in combination with an Intan RHD2132 miniature neural signal amplifier headstage PCB, recorded stimulation artifacts at a 20-kHz sampling rate, while a function generator (33220A, Keysight Technologies, Santa Rosa, CA) provided voltage pulses for LED driving. 50-ms long (5 Hz frequency, resulting in 25 % duty ratio) rectangular voltage pulses were used as LED drive signals. The off-time (low-level) voltage, on-time (high-level) voltage, pulse rise time, and pulse fall time were varied for different experiments. The experimental conditions used for each type of experiment are summarized in Supplementary Table 1. Before the LED drive signal was provided, the impedance (both the magnitude and the phase at 1 kHz) of each recording electrode on the µLED optoelectrode was measured using the Intan amplifier.

### Characterization of the effect of the shielding layer on the magnitude of *in vitro* LED-drive-induced artifact

Two one-metal-layer µLED optoelectrodes and two shielded µLED optoelectrodes, all of which were fabricated using the LED wafer with p^-^ silicon substrate (boron doped, N_A_ ≈ 5 × 10^16^ cm^-3^), were used. First, the high-level voltages required for generation of radiant flux of 230 nW - 11.5 µW (LED surface irradiance of 1.5 - 75 mW mm^-2^) were calculated. The high-level voltage of the LED drive pulse signal was varied according to the target irradiance, while the low-level voltage was fixed at 0 V and the rise time (as well as the fall time) was fixed as 5 ns (10 – 90 %, equivalent to 6.25 ns of 0 – 100 % rise and fall times).

### Characterization of the effect of the boron doping of the silicon substrate on the magnitude of *in vitro* LED-drive-induced artifact

Six shielded µLED optoelectrodes fabricated using LED wafers with FZ, p^-^, and p^+^ silicon substrate (two optoelectrodes from each wafer) were used. LED drive signals identical to those used for characterization of the effect of the shielding layer were used.

### Characterization of the effect of the transient stimulation pulse shaping on the magnitude of *in vitro* LED-drive-induced artifact

Two miniSTAR µLED optoelectrodes (shielded µLED optoelectrodes fabricated using LED wafers p^+^ silicon substrate) were used. The low-level voltage and the rise time of the LED drive pulse signal were varied, while the high-level voltage was fixed as 3.5 V. Low-level voltages of 0 V and 2.8 V were used, and rise and fall times (10 – 90 %) between 5 ns and 1 ms were used.

### Recording of *in vitro* LED-drive-induced artifact and data processing

For each experimental condition for each µLED, signals from the input channels of the neural signal amplifier IC were recorded for 30 seconds, so that artifact signals from longer than 100 pulses can be recorded. Average artifact signal was calculated by first highpass filtering the signal to remove low-noise fluctuations (with filters with 10 Hz and 250 Hz cutoff frequencies for wideband signal and highpass filtered signals, respectively) and calculating the average of the fifty 200-ms long segments in the middle of the 30 second period after the first 5 s of the recorded signal. Transient artifact magnitude was calculated from the difference between the maximum and the minimum values of high-pass filtered signal during the first 5-ms period from the point when the voltage changed from the off-level voltage. The mean transient artifact magnitude was calculated by taking the mean of the values from electrode whose impedance magnitudes are between 500 kΩ and 2 MΩ and the phases are between −80 ° and −55 ° at 1 kHz. Two µLED optoelectrodes from each cohort were used, and at least 21 electrodes per optoelectrode (out of 32 total, 25.83 in average) contributed to calculation of the mean artifact magnitude. The mean 1 kHz magnitude and phase of the electrode impedance of the electrodes which contributed to calculation of the mean artifact magnitude were 1.09 ± 0.09 MΩ and −68.2 ± 4.9 ° (mean ± SD, measured at 1 kHz).

### *in vivo* characterization and demonstrations of stimulation-artifact-free opto-electrophysiology

The animal procedures were approved by the Institutional Animal Care and Use Committee of the University of Michigan (protocol number PRO-7275). One male C57BL/6J mouse (32 g) was used for *in vivo* characterization. The mouse was kept on a regular 12 h - 12 h light - dark cycle and housed in pairs before surgery. No prior experimentation had been performed on this animal. Atropine (0.05 mg/kg, s.c.) was administered after isoflurane anesthesia induction to reduce saliva production. The body temperature was monitored and kept constant at 36 - 37 °C with a DC temperature controller (TCAT-LV; Physitemp, Clifton, NJ, USA). Stages of anesthesia were maintained by confirming the lack of nociceptive reflex. Skin of the head was shaved and the surface of the skull was cleaned by hydrogen peroxide (2 %). A 1-mm diameter craniotomy was drilled at 1.5 mm posterior from bregma and 2 mm lateral of the midline. The dura was removed over the dorsal CA1 region of the hippocampus and the mouse was injected with AAV5, CaMKII promoter driven ChR2 (AAV5-CaMKIIa-hChR2(H134R)-EYFP), resulting in expression of ChR2 in pyramidal neurons. Viruses were purchased from the University of North Carolina Vector Core (UNC-REF). After the surgery, the craniotomy was sealed with Kwik-Sil (World Precision Instruments, Sarasota, FL) until the day of recording.

On the day of recording, the mouse was anesthetized with isoflurane, the craniotomy was cleaned, and a shielded µLED optoelectrode with p^+^ silicon substrate was lowered to the CA1 region of the hippocampus. Baseline recording was performed (30 min), after which simultaneous recording and stimulation were done using three µLEDs from one shank (as described in Results in more details). 0.46 µW power, equivalent to 3 mW mm^-2^ irradiance at the surface of each µLED, was used to characterize the light induced artifact *in vivo* and to alter the activity of neurons (more details are provided in Results). For characterization of stimulation artifact and confirmation of optical induction of neuronal activities, pulsed optical stimulation (100-ms long, 2 Hz, 100 pulses) was generated from each µLED. The (10 - 90 %) rise and the fall times of each voltage pulse were set as 1 ms. After collecting sufficient data using optical stimulation from each µLED, a 500-ms long optical stimulation sequence involving switching on and off all the three µLEDs on the shank (whose details are provided in Results) were repeated 100 times. RHD2000 recording system with RHD2132 miniature neural signal amplifier headstage was used for acquisition of data from all the recording electrodes (n = 32, 20 kS/s sampling rate). Keysight 33220A function generator provided voltage pulses for LED driving.

A custom MATLAB (MathWorks, USA) script was used to calculate average stimulation artifact. Wideband traces were first high-pass filtered with a first-order filter with 250 Hz cutoff frequency to remove low-noise fluctuations. The average artifact signal from each recording channel was then obtained by averaging the middle 500-ms long segments (90 total segments out of 100).

The recorded data were then further analyzed for identification and clustering of action potentials. No manipulation in data (e.g. trimming of 1-ms long segments before and after the beginning and the ending of each pulsed optical stimulation) other than high-pass filtering (at 500 Hz) of the baseband signal was conducted. Spikes were first detected and automatically sorted using the Kilosort algorithm^35^ and then manually curated using Phy to get well-isolated single units (multi-unit and noise clusters were discarded). To measure the effect of LED stimulation on neuronal activity, peristimulus time histograms (PSTHs) were built around stimulus onset (spike trains were binned into 10-ms bins). Baseline and light-induced firing rate were calculated for each single unit, in which the baseline was defined as light-free epochs (400 ms) between trials and the stimulation period as the light-on (100 ms). Wilcoxon-signed rank test was used to compare the mean firing rate per trial (n = 100 trials) during baseline and LED stimulation.

## Acknowledgements

The work has been supported by NIH 1-U01-NS090526-01, NSF 1545858, NSF 1707316, NIH 1-R01-MH107396-01, and NIH 1-U01-NS090583-01.

## Author contributions

K. K. and E. Y. worked together on conceptual design of minimal-stimulation-artifact (miniSTAR) µLED optoelectrodes. K. K. designed, fabricated, and assembled µLED optoelectrodes; conducted CAD simulations; conducted *in vitro* characterizations; and analyzed recorded data. M. V. conducted *in vivo* experiments with K. K. and processed recorded neuronal signals. J. P. S., K. D. W., G. B. and E. Y. supervised the study. K.K., M. V., and E. Y. wrote the manuscript. All authors discussed the results and commented and edited the manuscript.

