## Supplementary Information for "Artifact-free, high-temporal-resolution *in vivo* opto-electrophysiology with microLED optoelectrodes"

**for**

**Supplementary Results**

**Optical power and efficiency distributions of μLEDs fabricated on GaN-on-Si LED wafers with differently boron doped silicon substrates.**

Electrical and optical characteristics of μLEDs fabricated on GaN-on-Si GaN/InGaN MQW LED wafers with different doping densities were measured and compared. After current and radiant flux values were measured at varying LED bias voltages from 2 V to 4 V, current at forward bias voltage of 4 V (I(V = 4 V)), radiant flux generated from forward bias of 4 V ($E_e$(V = 4 V)), and the peak plug efficiency (max($\eta_{plug}$)) of each LED were calculated and compared.

Contrary to our initial conjecture that, due to increased defect density inside silicon substrate, the efficiency and therefore the maximum output radiant flux of LEDs on heavily boron doped silicon substrate will be lower than those of the LEDs on intrinsic silicon substrate, the efficiency of the LEDs on the intrinsic silicon substrate were the lowest (Supplementary Figure 1a-c). However, considering large variation of LED characteristics within a wafer (Supplementary Figure 1d-f), it is difficult to conclude that the doping density of the silicon substrate has significant effect on the electrical and the optical characteristics of the LEDs and therefore their efficiency.

**Simulation of electric field intensity inside μLED optoelectrode shanks during optical stimulation**

We built models of one-metal-layer and shielded μLED optoelectrode shanks and calculated mutual capacitances between the interconnects (Supplementary Figure 2c and 2d, top). Because n-type gallium nitride (n-GaN) layer underneath the interconnects is also a part of the LED drive circuitry and its electrostatic potential changes as a function of LED forward bias voltage, the layer was also taken into account as an electrode in the model.

In the one-metal-layer μLED optoelectrode structure, the capacitance between an LED interconnect and a recording electrode interconnect that are the closest with each other was calculated to be 5.37 × $10^{-19}$ F $\mu m^{-1}$, and that between the n-GaN layer and one recording electrode interconnect was 2.3 × $10^{-16}$ F $\mu m^{-1}$. When all the interconnects were assumed to be floating at both ends, capacitive voltage coupling between the LED interconnects and the recording electrode interconnect was -48.96 dB (3.57 mV coupling for 1 V LED voltage), and the coupling between the n-GaN layer and the recording electrode interconnects was -0.06 dB (0.99 mV coupling for 1 mV n-GaN voltage). Voltage distribution inside the one-metal-layer optoelectrode shank (and the air surrounding the optoelectrode) is shown in Supplementary Figure 2c (bottom).

Electrostatic simulation of shielded μLED optoelectrode structure predicted that the coupling would be greatly reduced (Supplementary Figure 2d, bottom). Reduction of coupling between LED interconnects recording interconnects was greater than 46 orders of magnitude (to approximately - 975 dB). Coupling between the n-GaN layer and the recording interconnects was reduced by 8 orders of magnitude (to approximately - 60 dB). It should be noted that the simulation expected ideal ground-connected shielding layer and floating electrodes and therefore the values seem greatly exaggerated.

**Simulation of PV-induced voltage around μLED optoelectrode shanks during optical stimluation**

We built a 3D model of a μLED optoelectrode shank and simulated the effect of illumination on silicon substrate (Supplementary Figure 5a). The doping density of boron, an acceptor dopant, inside the silicon substrate and the intensity of the optical illumination were varied. We observed a series of phenomena that results in buildup of electrostatic potential at the substrate-electrolyte interface and in turn generation of a voltage pulse with negative polarity in the recorded signal (Supplementary Figure 5b). First, optical illumination induced electron-hole pair generation inside silicon substrate, and optically generated carriers redistributed inside the substrate separately depending on their types. Difference between electron and hole distribution patterns gave rise to electric field inside the substrate, and, in turn, the electrostatic potential of the substrate-electrolyte interface changed.

Because the electrolyte is connected to the common reference pin of the amplifier chip which is then connected to the inverting inputs of the low-noise amplifiers in the IC, the resulting output waveform would have a negative polarity.

Supplementary Figure 5c shows calculated substrate-electrolyte interface electrostatic potential (voltage) for substrates with different doping densities under illumination with different intensities. It is worth noting that, while higher doping density resulted in lower voltage at lower irradiance, with higher irradiance, the voltage on lightly doped (typically referred to as $p^-$) substrates became higher than that on substrates that are almost intrinsic (not doped, typically referred to as HR or FZ, especially if the silicon substrate was float-zone grown for high-purity and low doping density). As can be seen in Supplementary Figure 5d, it was calculated that the interface voltage from a substrate with boron doping density of $5 \times 10^{16}$ $cm^{-3}$ ($p^-$ substrate) can be as high as that from the substrate with boron doping density of $4 \times 10^{12}$ $cm^{-3}$ (FZ substrate) under illumination with irradiance as high as 50 mW $mm^{-2}$. On the other hand, the interface voltage from the substrate with boron doping density of $1 \times 10^{20}$ $cm^{-3}$ ($p^+$ substrate) was kept relatively low even with high intensity illumination.

**Supplementary Methods**

**Characterization of optical power and efficiency distributions of μLEDs fabricated on wafers with Si substrates with different doping densities**

Simple μLED test structures were built using the process similar to fabrication process for μLED optoelectrodes. After formation of μLED structures, whose LED mesa dimensions are identical to those on the μLED optoelectrodes (23 × 10 μm, only 15 × 10 μm of which is exposed on the front side), GaN-on-Si GaN/InGaN MQW LED wafers with μLED test structures were diced into small (4 × 10 mm) pieces each of which contains nine μLEDs. The pieces were then mounted on the PCBs and connected in the same way as the actual μLED optoelectrodes are connected to the PCBs.

The electrical and optical characteristics of each μLED on the μLED test structures were characterized using the setup and the procedure identical to those for characterization of the actual μLED optoelectrodes, outlined in Methods.

**Electrostatics simulation for calculation of mutual capacitances between interconnects**

COMSOL Multiphysics (COMSOL Inc., Burlington, MA) was used for finite element analysis of mutual capacitance distribution among the metal traces (interconnects) on the shanks of the μLED optoelectrode. A 2D model of the shank was built by drawing the cross-section of the optoelectronic shank, and the electrostatics physics interface was imported to calculate the mutual capacitance values of 100-μm long segments of the of the shanks of μLED optoelectrodes with and without shielding layers. Built-in material properties (dielectric constants) of air and silicon dioxide were used. Each interconnect plus the n-GaN layer was assigned either terminal (V = 0) or floating potential (Q = 0) boundary condition. Automatic terminal sweep was used for calculation of Maxwell capacitance matrix. Mutual capacitance values were then extracted from the matrix. For calculation of capacitive voltage coupling magnitude, all the boundaries that correspond to recording electrode interconnects were defined as terminals with floating potential, all the boundaries that correspond to LED ground (cathode) interconnects and shielding layer were defined as grounds (V = 0), and then, assuming the LED and n-GaN voltages of 1 V, the voltage values (in dB) were reported.

#### ***In vitro* characterization of photovoltaic voltage induction on μLED optoelectrodes and electrode arrays on non-Si substrates**

Identical to characterization of LED-drive-induced stimulation artifact, characterization of photovoltaic voltage induction was conducted in 1 × PBS solution (prepared using 10 × PSB purchased from MP Biomedicals, Solon, OH). A fiber optic cannula (CFMXD10, Thorlabs, Newton, NJ) was attached to a clear plastic container (Container Store, Coppel, TX) through a hole drilled on a side of the container using a 3D printed frame and glue. PBS was poured into the container until the exposed optical fiber tip of the cannula is submerged approximately 2.5 mm under the surface of PBS. The optoelectrode, attached to a 3-axis micromanipulator on a stereotaxic frame (Model 962, David Kopf Instruments, Tujunga, CA), was lowered into the container until its shanks were sufficiently submerged into the PBS. The position of the optoelectrode was precisely adjusted using the micromanipulator so that the tips of the optoelectrode are exactly 1.5 mm away from the tip of the optical fiber, while the top side of the optoelectrode which has the electrodes and the μLED are facing the optical fiber.

Optical stimulation was provided using a fiber-coupled LED light source (M470F3, Thorlabs), whose spectrum ($\lambda_{peak}$ = 470 nm) is similar to those of the μLEDs on the μLED optoelectrodes. Optical power at the end of the fiber optic cannula was measured beforehand using the combination of the integrating sphere and the spectrometer. 5-Hz, 50-ms long (25 % duty ratio) rectangular voltage pulses with varying on-voltage levels were used as the LED drive signal. Pulses with 0 V low-level (off-time) voltage and high-level (on-time) voltage that would generate the irradiance same as the pre-characterized irradiance was used.

Intan RHD2132 neural signal amplifier headstage PCB (RHD2132, Intan Technologies, Los Angeles, CA) recorded the induced voltage signal. Data collection and processing followed the procedure identical to that has been previously outlined in Methods for LED-drive-induced stimulation artifact characterization.

**Device physics simulation for calculation of photovoltaic-effect-induced electrostatic potential buildup inside silicon substrate**

Sentaurus TCAD suite (Synopsis, Mountain View, CA) was used for finite element analysis of carrier generation and electrostatic potential buildup inside μLED optoelectrode's silicon substrate during LED illumination. A 3D model of a shank was built using Sentaurus Device Editor and the shank's silicon substrate was given variable boron doping density. Two contacts, each of which indicate GaN-AlN interface and the silicon-PBS interface, were created and appropriately assigned. Using Sentaurus Device, carrier distribution and electrostatic potential buildup during irradiation of specified intensity were calculated. Ground ($V(t) = 0$) and floating ($Q(t) = Q_{0,\ stationary}$) boundary conditions were applied to the ground and the electrolyte contacts, respectively, before simulation. First, steady-state conditions were calculated. For time domain simulation, a 23-μm wide and 10-μm long rectangular region under the AlN buffer was defined as the light source. Coupled Poisson equations for electrons and holes were then solved in time domain with assumption of density-dependent Shockley-Read-Hall recombination. For each iteration, different irradiance value was used, while the wavelength of the light was kept at 470 nm. Voltage of the silicon-PBS interface before, during, and after specified irradiation and optical generation of electrons and holes was recorded and reported for each combination of boron doping density and irradiance.

**Measurement of stimulation artifact during current-based μLED driving**

Setup for *in vitro* characterization of stimulation artifact, described in Methods, was used. Two miniSTAR μLED optoelectrodes, identical to those used for characterization of stimulation artifact resulting from voltage driving, were used. Instead of a Keysight 33220A function generator, a custom FPGA-controlled current source (DAC8750, Texas Instruments, Dallas, TX) was used as the LED driver. Current pulses with different rise times and rising and falling edge shapes were generated using SPI commands created with a custom script written in Matlab (Mathworks, Natick, MA). Low- and high-level values of the current pulses were kept as 0 A and 75 μA regardless of the type of waveform.

**Detailed shielded μLED optoelectrode fabrication process**

All microfabrication steps were conducted in the Lurie Nanofabrication Facility (LNF) of the University of Michigan, Ann Arbor, MI. For all photolithography processes, MEGAPOSIT SPR 220 3.0 and 7.0 (Dow Electronic Materials, DuPont de Nemours, Inc., Wilmington, DE) photoresists were used in combination with an i-line stepper (GCA AS200 AutoStep, General Signal Corp., Stamford, CT) for exposure.

First, LED wafers were cleaned in acetone, isopropyl alcohol (IPA), and then deionized wafer (DI $H_2O$) for removal of organic residue. Next, the wafers were exposed to hydrochloric acid (HCl) for surface cleaning. Following wafer surface cleaning, chlorine-based reactive ion etching (RIE) was used for LED mesa definition (using LAM 9400, Lam Research Corporation, Fremont, CA). Plasma-enhanced chemical vapor deposition (PECVD) of silicon dioxide ($SiO_2$) was used for surface passivation of LED structure (using GSI ULTRADEP 2000, GSI Lumonics, Novanta Inc., Bedford, MA, USA). After $SiO_2$ PECVD, $SiO_2$ on top of the p- and n-GaN contact sites were etched using $C_4F_8$/$SF_6$-based RIE (using LAM 9400). Lift-off patterned, electron-beam (e-beam) evaporated nickel/gold (Ni/Au, deposited using SJ-20, Denton Vacuum, Moorestown, NJ) and titanium/aluminum/titanium/gold (Ti/Al/Ti/Au, deposited using Enerjet Evaporator, K. J. Lesker, Jefferson Hills, PA and Denton Vacuum) metal stacks were used for p-

GaN contact and n-GaN contact, respectively. Following deposition, p-GaN contact metal stack was annealed at 500 °C in $N_2/O_2$ environment (using JetFirst 150, SEMCO Inc., Irving, TX).

Bilayer stack of atomic-layer-deposited (ALD) aluminum oxide ($Al_2O_3$, deposited using Oxford OpAL, Oxford Instruments, Abingdon, UK) and PECVD $SiO_2$ (deposited using P5000 PECVD, Applied Materials, Santa Clara, CA) composed the passivation layers, and e-beam evaporated Ti/Au (deposited using Enerjet Evaporator) formed the metal layers. After deposition of ALD $Al_2O_3$ and PECVD $SiO_2$ bilayer above the top metal layer, the top passivation was partially etched etching using combination of RIE (using LAM 9400) and wet etching (using dilute buffered hydrofluoric acid) to expose recording electrode contact sites. Recording electrodes were defined using lift-off patterning of sputter-deposited titanium/platinum/iridium (Ti/Pt/Ir, deposited using LAB 18, K. J. Lesker) stack.

Double-sided plasma dicing process, consisting of patterned front-side deep reactive ion etching (DRIE) followed by backside plasma thinning, was conducted (using STS Pegasus 4, SPTS Technologies, Orbotech Ltd., Yavne, Israel). Released optoelectrodes were cleaned in xylenes, acetone, and then IPA before pick up and assembly.

**Supplementary Tables**

**Supplementary Table 1: Summary of the experimental conditions used for each type of experiment for characterization of stimulation artifact.**

| **Type of experiment** | | **Effect of shielding layer** | **Effect of substrate doping** | **Effect of transient pulse shaping** |
|---|---|---|---|---|
| **Devices used** | | 2 × one-metal-layer, $p^-$-Si substrate;<br>2 × shielded, $p^-$-Si substrate | 2 × shielded, FZ-Si substrate;<br>2 × shielded, $p^-$-Si substrate;<br>2 × shielded, $p^+$-Si substrate | 2 × shielded, $p^+$-Si substrate (miniSTAR) |
| **LED signal** | **Low-level voltage (V)** | 0 | 0 | 0, 2.8 |
| | **High-level voltage (V)** | [2.71 ± 0.02, 3.36 ± 0.09] (mean ± SD) | [2.76 ± 0.11, 3.57 ± 0.15] (mean ± SD) | 3.5 |
| | **Equivalent on-time radiant flux (μW)** | 0.23, 0.46, 1.15, 2.3, 4.6, 6.9, 9.2, 11.5 | 0.23, 0.46, 1.15, 2.3, 4.6, 6.9, 9.2, 11.5 | 9.29 ± 2.47 (mean ± SD) |
| | **Pulse rise time (10 - 90 %)** | 5 ns | 5 ns | 5 ns, 10 ns, 50 ns, 100 ns, 500 ns, 1 μs, 5 μs, 10 μs, 50 μs, 100 μs, 500 μs, 1 ms |

**Supplementary Table 2: List of the statistical tests of significance used in the study**.

| | |
|---|---|
| **Figure (panel)** | **Fig. 2b** |
| **Test used** | Mann-Whitney U test with Bonferroni correction |
| **Samples and categories** | Peak-to-peak magnitudes of stimulation artifact, recorded from all sites on the one-metal-layer (non-shielded) μLED optoelectrodes, measured during optical stimulation using μLEDs resulting in LED surface irradiance of 1.5, 3, 7.5, 15, 30, 45, 60, and 75 mW $mm^{-2}$ |
| **Statistics provided in figure** | Box plots with whiskers and outliers (denoting median, IQR, EVs and outliers) |
| **n** | 75 (for all categories) |
| **p-values** | $2.84 \times 10^{-1}$ (1.5 vs. 3 mW $mm^{-2}$), $3.25 \times 10^{-2}$ (3 vs. 7.5 mW $mm^{-2}$), $3.89 \times 10^{-3}$ (7.5 vs. 15 mW $mm^{-2}$), $7.87 \times 10^{-3}$ (15 vs. 30 mW $mm^{-2}$), $1.63 \times 10^{-2}$ (30 vs. 45 mW $mm^{-2}$), $3.61 \times 10^{-1}$ (45 vs. 60 mW $mm^{-2}$), & $2.48 \times 10^{-3}$ (60 vs. 75 mW $mm^{-2}$) |
| **Other values** | z = -1.07 (1.5 vs. 3 mW $mm^{-2}$), 2.14 (3 vs. 7.5 mW $mm^{-2}$), -2.89 (7.5 vs. 15 mW $mm^{-2}$), 2.66 (15 vs. 30 mW $mm^{-2}$), -2.40 (30 vs. 45 mW $mm^{-2}$), $9.13 \times 10^{-1}$ (45 vs. 60 mW $mm^{-2}$), & -3.03 (60 vs. 75 mW $mm^{-2}$) |

| | |
|---|---|
| **Figure (panel)** | **Fig. 2e** |
| **Test used** | Mann-Whitney U test with Bonferroni correction |
| **Samples and categories** | Peak-to-peak magnitudes of stimulation artifact, recorded from all sites on shielded μLED optoelectrodes fabricated using LED wafer with moderately boron-doped silicon substrate, measured during optical stimulation using μLEDs resulting in in LED surface irradiance of 1.5, 3, 7.5, 15, 30, 45, 60, and 75 mW $mm^{-2}$ |
| **Statistics provided in figure** | Box plots with whiskers and outliers (denoting median, IQR, EVs and outliers) |
| **n** | 67 (for all categories) |
| **p-values** | $1.76 \times 10^{-1}$ (1.5 vs. 3 mW $mm^{-2}$), $9.70 \times 10^{-2}$ (3 vs. 7.5 mW $mm^{-2}$), $4.03 \times 10^{-6}$ (7.5 vs. 15 mW $mm^{-2}$), $3.51 \times 10^{-10}$ (15 vs. 30 mW $mm^{-2}$), $1.88 \times 10^{-7}$ (30 vs. 45 mW $mm^{-2}$), $2.23 \times 10^{-5}$ (45 vs. 60 mW $mm^{-2}$), & $2.48 \times 10^{-3}$ (60 vs. 75 mW $mm^{-2}$) |
| **Other values** | z = 1.35 (1.5 vs. 3 mW $mm^{-2}$), -1.66 (3 vs. 7.5 mW $mm^{-2}$), -4.61 (7.5 vs. 15 mW $mm^{-2}$), -6.27 (15 vs. 30 mW $mm^{-2}$), -5.21 (30 vs. 45 mW $mm^{-2}$), -4.24 (45 vs. 60 mW $mm^{-2}$), & -3.03 (60 vs. 75 mW $mm^{-2}$) |

| Figure (panel) | Fig. 2g |
|---|---|
| Test used | Mann-Whitney U test |
| Samples and categories | Peak-to-peak magnitudes of stimulation artifact, recorded from all sites on the 1$^{st}$-generation (i.e. non-shielded with moderate boron doping of the silicon substrate) $\mu$LED optoelectrodes and shielded $\mu$LED optoelectrodes fabricated using LED wafer with moderately boron-doped silicon substrate, measured during optical stimulation using $\mu$LEDs resulting in LED surface irradiance of 1.5, 3, 7.5, 15, 30, 45, 60, and 75 mW mm$^{-2}$ |
| Statistics provided in figure | Scatter plots with error bars (denoting mean and SD) |
| n | 75 and 67 (for all categories) |
| p-values | $4.09 \times 10^{-23}$ (@ 1.5 mW mm$^{-2}$), $3.07 \times 10^{-23}$ (@ 3 mW mm$^{-2}$), $9.70 \times 10^{-24}$ (@ 7.5 mW mm$^{-2}$), $5.20 \times 10^{-24}$ (@ 15 mW mm$^{-2}$), $2.18 \times 10^{-17}$ (@ 30 mW mm$^{-2}$), $8.72 \times 10^{-22}$ (@ 45 mW mm$^{-2}$), $2.81 \times 10^{-15}$ (@ 60 mW mm$^{-2}$), & $2.72 \times 10^{-23}$ (@ 75 mW mm$^{-2}$) |
| Other values | z = 9.90 (@ 1.5 mW mm$^{-2}$), 9.93 (@ 3 mW mm$^{-2}$), $1.00 \times 10^{1}$ (@ 7.5 mW mm$^{-2}$), 8.48 (@ 15 mW mm$^{-2}$), 9.59 (@ 30 mW mm$^{-2}$), 7.90 (@ 45 mW mm$^{-2}$), 7.90 (@ 60 mW mm$^{-2}$), & 9.94 (@ 75 mW mm$^{-2}$) |

| Figure (panel) | Fig. 3a, left |
|---|---|
| Test used | Mann-Whitney U test with Bonferroni correction |
| Samples and categories | Peak-to-peak magnitudes of stimulation artifact, recorded from all sites on shielded $\mu$LED optoelectrodes fabricated using LED wafer with FZ-silicon substrate, measured during optical stimulation using $\mu$LEDs resulting in LED surface irradiance of 1.5, 3, 7.5, 15, 30, 45, 60, and 75 mW mm$^{-2}$ |
| Statistics provided in figure | Box plots with whiskers and outliers (denoting median, IQR, EVs and outliers) |
| n | 124 (for all categories) |
| p-values | $1.98 \times 10^{-15}$ (1.5 vs. 3 mW mm$^{-2}$), $4.91 \times 10^{-28}$ (3 vs. 7.5 mW mm$^{-2}$), $5.67 \times 10^{-13}$ (7.5 vs. 15 mW mm$^{-2}$), $2.81 \times 10^{-8}$ (15 vs. 30 mW mm$^{-2}$), $8.95 \times 10^{-3}$ (30 vs. 45 mW mm$^{-2}$), $5.29 \times 10^{-2}$ (45 vs. 60 mW mm$^{-2}$), & $9.61 \times 10^{-1}$ (60 vs. 75 mW mm$^{-2}$) |
| Other values | z = -7.94 (1.5 vs. 3 mW mm$^{-2}$), $-1.10 \times 10^{1}$ (3 vs. 7.5 mW mm$^{-2}$), -7.21 (7.5 vs. 15 mW mm$^{-2}$), -5.55 (15 vs. 30 mW mm$^{-2}$), -2.61 (30 vs. 45 mW mm$^{-2}$), -1.94 (45 vs. 60 mW mm$^{-2}$), & $-4.87 \times 10^{-2}$ (60 vs. 75 mW mm$^{-2}$) |

| Figure (panel) | Fig. 3a, center |
|---|---|
| **Test used** | Mann-Whitney U test with Bonferroni correction |
| **Samples and categories** | Peak-to-peak magnitudes of stimulation artifact, recorded from all sites on shielded μLED optoelectrodes fabricated using LED wafer with moderately boron-doped silicon substrate, measured during optical stimulation using μLEDs resulting in LED surface irradiance of 1.5, 3, 7.5, 15, 30, 45, 60, and 75 mW $mm^{-2}$ |
| **Statistics provided in figure** | Box plots with whiskers and outliers (denoting median, IQR, EVs and outliers) |
| **n** | 67 (for all categories) |
| **p-values** | $1.76 \times 10^{-1}$ (1.5 vs. 3 mW $mm^{-2}$), $9.70 \times 10^{-2}$ (3 vs. 7.5 mW $mm^{-2}$), $4.03 \times 10^{-6}$ (7.5 vs. 15 mW $mm^{-2}$), $3.51 \times 10^{-10}$ (30 vs. 45 mW $mm^{-2}$), $1.88 \times 10^{-7}$ (30 vs. 45 mW $mm^{-2}$), $2.23 \times 10^{-5}$ (45 vs. 60 mW $mm^{-2}$), & $2.48 \times 10^{-3}$ (60 vs. 75 mW $mm^{-2}$) |
| **Other values** | z = 1.35 (1.5 vs. 3 mW $mm^{-2}$), -1.66 (3 vs. 7.5 mW $mm^{-2}$), -4.61 (7.5 vs. 15 mW $mm^{-2}$), -6.27 (15 vs. 30 mW $mm^{-2}$), -5.21 (30 vs. 45 mW $mm^{-2}$), -4.24 (45 vs. 60 mW $mm^{-2}$), & -3.03 (60 vs. 75 mW $mm^{-2}$) |

| Figure (panel) | Fig. 3a, right |
|---|---|
| **Test used** | Mann-Whitney U test with Bonferroni correction |
| **Samples and categories** | Peak-to-peak magnitudes of stimulation artifact, recorded from all sites on shielded μLED optoelectrodes fabricated using LED wafer with heavily boron-doped silicon substrate, measured during optical stimulation using μLEDs resulting in LED surface irradiance of 1.5, 3, 7.5, 15, 30, 45, 60, and 75 mW $mm^{-2}$ |
| **Statistics provided in figure** | Box plots with whiskers and outliers (denoting median, IQR, EVs and outliers) |
| **n** | 151 (for all categories) |
| **p-values** | $9.61 \times 10^{-1}$ (1.5 vs. 3 mW $mm^{-2}$), $5.22 \times 10^{-1}$ (3 vs. 7.5 mW $mm^{-2}$), $6.05 \times 10^{-1}$ (7.5 vs. 15 mW $mm^{-2}$), $8.68 \times 10^{-1}$ (15 vs. 30 mW $mm^{-2}$), $6.46 \times 10^{-1}$ (30 vs. 45 mW $mm^{-2}$), $4.78 \times 10^{-1}$ (45 vs. 60 mW $mm^{-2}$), & $5.54 \times 10^{-1}$ (60 vs. 75 mW $mm^{-2}$) |
| **Other values** | z = $4.88 \times 10^{-2}$ (1.5 vs. 3 mW $mm^{-2}$), $6.41 \times 10^{-1}$ (3 vs. 7.5 mW $mm^{-2}$), $5.17 \times 10^{-1}$ (7.5 vs. 15 mW $mm^{-2}$), $-1.66 \times 10^{-1}$ (15 vs. 30 mW $mm^{-2}$), $-4.60 \times 10^{-1}$ (30 vs. 45 mW $mm^{-2}$), $-7.09 \times 10^{-1}$ (45 vs. 60 mW $mm^{-2}$), & $-5.92 \times 10^{-1}$ (60 vs. 75 mW $mm^{-2}$) |

| Figure (panel) | Fig. 4b, left |
|---|---|
| Test used | Mann-Whitney U test with Bonferroni correction |
| Samples and categories | Peak-to-peak magnitudes of stimulation artifact, recorded from all sites on shielded μLED optoelectrodes fabricated using LED wafer with FZ-silicon substrate (FZ-Si), shielded μLED optoelectrodes fabricated using LED wafer with moderately boron-doped silicon substrate ($p^-$-Si), and shielded μLED optoelectrodes fabricated using LED wafer with heavily boron-doped silicon substrate ($p^+$-Si) |
| Statistics provided in figure | Box plots with whiskers and outliers (denoting median, IQR, EVs and outliers) |
| n | 124 (FZ-Si), 67 ($p^-$-Si), & 151 ($p^+$-Si) |
| p-values | $1.30 \times 10^{-5}$ ($p^-$-Si vs. FZ-Si) & $5.64 \times 10^{-24}$ ($p^-$-Si vs. $p^+$-Si) |
| Other values | z = -4.36 ($p^-$-Si vs. FZ-Si) & 100.98 ($p^-$-Si vs. $p^+$-Si) |

| Figure (panel) | Fig. 4b, center |
|---|---|
| Test used | Mann-Whitney U test with Bonferroni correction |
| Samples and categories | Peak-to-peak magnitudes of stimulation artifact, recorded from two sites at the bottom of each shank (sites 1 & 2) on shielded μLED optoelectrodes fabricated using LED wafer with FZ-silicon substrate (FZ-Si), shielded μLED optoelectrodes fabricated using LED wafer with moderately boron-doped silicon substrate ($p^-$-Si), and shielded μLED optoelectrodes fabricated using LED wafer with heavily boron-doped silicon substrate ($p^+$-Si) |
| Statistics provided in figure | Box plots with whiskers and outliers (denoting median, IQR, EVs and outliers) |
| n | 34 (FZ-Si), 19 ($p^-$-Si), & 38 ($p^+$-Si) |
| p-values | $1.19 \times 10^{-2}$ ($p^-$-Si vs. FZ-Si) & $9.53 \times 10^{-1}$ ($p^-$-Si vs. $p^+$-Si) |
| Other values | z = -2.51 ($p^-$-Si vs. FZ-Si) & $4.92 \times 10^{-2}$ ($p^-$-Si vs. $p^+$-Si) |

| Figure (panel) | Fig. 4b, right |
|---|---|
| Test used | Mann-Whitney U test with Bonferroni correction |
| Samples and categories | Peak-to-peak magnitudes of stimulation artifact, recorded from two sites at the top of each shank (sites 7 & 8) on shielded μLED optoelectrodes fabricated using LED wafer with FZ-silicon substrate (FZ-Si), shielded μLED optoelectrodes fabricated using LED wafer with moderately boron-doped silicon substrate ($p^-$-Si), and shielded μLED optoelectrodes fabricated using LED wafer with heavily boron-doped silicon substrate ($p^+$-Si) |
| Statistics provided in figure | Box plots with whiskers and outliers (denoting median, IQR, EVs and outliers) |
| n | 32 (FZ-Si), 22 ($p^-$-Si), & 41 ($p^+$-Si) |
| p-values | $1.32 \times 10^{-5}$ ($p^-$-Si vs. FZ-Si) & $8.29 \times 10^{-11}$ ($p^-$-Si vs. $p^+$-Si) |
| Other values | z = -4.36 ($p^-$-Si vs. FZ-Si) & 6.50 ($p^-$-Si vs. $p^+$-Si) |

| Figure (panel) | Fig. 4e |
|---|---|
| **Test used** | Mann-Whitney U test |
| **Samples and categories** | Peak-to-peak magnitudes of stimulation artifact, recorded from different sites on each shank (sites 1 - 8) on shielded μLED optoelectrodes fabricated using LED wafer with heavily boron-doped silicon substrate; comparing two sites with same LED-to-interconnect distances |
| **Statistics provided in figure** | Box plots with whiskers and outliers (denoting median, IQR, EVs and outliers) |
| **n** | 22 (site 1), 16 (site 2), 12 (site 3), 22 (site 4), 20 (site 5), 18 (site 6), 22 (site 7), & 19 (site 8) |
| **p-values** | $7.79 \times 10^{-1}$ (site 1 vs. 2), $3.97 \times 10^{-1}$ (site 3 vs. 4), $5.93 \times 10^{-2}$ (site 5 vs. 6), & $5.05 \times 10^{-1}$ (site 7 vs. 8) |
| **Other values** | $z = 2.81 \times 10^{-1}$ (site 1 vs. 2), $8.47 \times 10^{-1}$ (site 3 vs. 4), 1.89 (site 5 vs. 6), & $-6.67 \times 10^{-1}$ (site 7 vs. 8) |

| Figure (panel) | Fig. 4e |
|---|---|
| **Test used** | Mann-Whitney U test with Bonferroni correction |
| **Samples and categories** | Peak-to-peak magnitudes of stimulation artifact, recorded from different sites on each shank (sites 1 - 8) on shielded μLED optoelectrodes fabricated using LED wafer with heavily boron-doped silicon substrate; comparing pairs of two sites with same LED-to-interconnect distances with each other pair |
| **Statistics provided in figure** | Box plots with whiskers and outliers (denoting median, IQR, EVs and outliers) |
| **n** | 38 (site 1 & 2), 44 (site 3 & 4), 38 (site 5 & 6), & 41 (site 7 & 8) |
| **p-values** | $4.65 \times 10^{-11}$ (site 1 & 2 vs. sites 3 & 4), $1.41 \times 10^{-4}$ (site 3 & 4 vs. sites 5 & 6), & $6.99 \times 10^{-4}$ (site 5 & 6 vs. sites 7 & 8) |
| **Other values** | z = 6.58 (site 1 & 2 vs. sites 3 & 4), 3.81 (site 3 & 4 vs. sites 5 & 6), & 3.39 (site 5 & 6 vs. sites 7 & 8) |

| Figure (panel) | Fig. 5d |
|---|---|
| **Test used** | Mann-Whitney U test with Bonferroni correction |
| **Samples and categories** | Peak-to-peak magnitudes of stimulation artifact, recorded from two sites at the bottom of each shank (sites 1 & 2) on μLTAR optoelectrodes during LED driving with voltage pulses with 0 V low-level voltage and 5 ns rise time (0V-5ns), with pulses with 0 V low-level voltage and 1 ms rise time (0V-1ms), with pulses with 2.8 V low-level voltage and 5 ns rise time (2P8V-4ns), and with pulses with 2.8V low-level voltage and 1 ms rise time (2P8V-1ms) |
| **Statistics provided in figure** | Box plots with whiskers (denoting median, IQR, and EVs) |
| **n** | 35 (for all categories) |
| **p-values** | $1.98 \times 10^{-12}$ (0V-5ns vs. 0V-1ms), $6.55 \times 10^{-13}$ (0V-5ns vs. 2P8V-5ns), & $6.55 \times 10^{-13}$ (0V-5ns vs. 2P8V-1ms) |
| **Other values** | z = 7.04 (0V-5ns vs. 0V-1ms), 7.19 (0V-5ns vs. 2P8V-5ns), & 7.19 (0V-5ns vs. 2P8V-1ms) |

| Figure (panel) | Supplementary Figure 1a, bottom |
|---|---|
| **Test used** | Mann-Whitney U test with Bonferroni correction |
| **Samples and categories** | Current through μLEDs fabricated on LED wafer with FZ-silicon substrate (FZ-Si), μLEDs fabricated on LED wafer with moderately boron-doped silicon substrate ($p^-$-Si), and μLEDs fabricated on LED wafer with heavily boron-doped silicon substrate ($p^+$-Si), at 4 V of forward bias voltage |
| **Statistics provided in figure** | Box plots with whiskers and outliers (denoting median, IQR, EVs and outliers) |
| **n** | 43 (FZ-Si), 44 ($p^-$-Si), & 43 ($p^+$-Si) |
| **p-values** | $3.01 \times 10^{-5}$ (FZ-Si vs. $p^-$-Si), $1.34 \times 10^{-5}$ (FZ-Si vs. $p^+$-Si), & $2.68 \times 10^{-1}$ ($p^-$-Si vs. $p^+$-Si) |
| **Other values** | z = -4.17 (FZ-Si vs. $p^-$-Si), -4.35 (FZ-Si vs. $p^+$-Si), & -1.11 ($p^-$-Si vs. $p^+$-Si) |

| Figure (panel) | Supplementary Figure 1b, bottom |
|---|---|
| **Test used** | Mann-Whitney U test with Bonferroni correction |
| **Samples and categories** | Radiant flux generated from μLEDs fabricated on LED wafer with FZ-silicon substrate (FZ-Si), μLEDs fabricated on LED wafer with moderately boron-doped silicon substrate ($p^-$-Si), and μLEDs fabricated on LED wafer with heavily boron-doped silicon substrate ($p^+$-Si), at 4 V of forward bias voltage |
| **Statistics provided in figure** | Box plots with whiskers and outliers (denoting median, IQR, EVs and outliers) |
| **n** | 43 (FZ-Si), 44 ($p^-$-Si), & 43 ($p^+$-Si) |
| **p-values** | $7.52 \times 10^{-6}$ (FZ-Si vs. $p^-$-Si), $2.62 \times 10^{-6}$ (FZ-Si vs. $p^+$-Si), & $3.35 \times 10^{-1}$ ($p^-$-Si vs. $p^+$-Si) |
| **Other values** | z = -4.48 (FZ-Si vs. $p^-$-Si), -4.70 (FZ-Si vs. $p^+$-Si), & $-9.64 \times 10^{-1}$ ($p^-$-Si vs. $p^+$-Si) |

| Figure (panel) | Supplementary Fig. 1c, bottom |
|---|---|
| Test used | Mann-Whitney U test with Bonferroni correction |
| Samples and categories | Maximum plug efficiency of μLEDs fabricated on LED wafer with FZ-silicon substrate (FZ-Si), μLEDs fabricated on LED wafer with moderately boron-doped silicon substrate ($p^-$-Si), and μLEDs fabricated on LED wafer with heavily boron-doped silicon substrate ($p^+$-Si) |
| Statistics provided in figure | Box plots with whiskers and outliers (denoting median, IQR, EVs and outliers) |
| n | 43 (FZ-Si), 44 ($p^-$-Si), & 43 ($p^+$-Si) |
| p-values | $1.10 \times 10^{-2}$ (FZ-Si vs. $p^-$-Si), $8.57 \times 10^{-2}$ (FZ-Si vs. $p^+$-Si), & $4.52 \times 10^{-1}$ ($p^-$-Si vs. $p^+$-Si) |
| Other values | z = -2.54 (FZ-Si vs. $p^-$-Si), -1.72 (FZ-Si vs. $p^+$-Si), & $-7.51 \times 10^{-1}$ ($p^-$-Si vs. $p^+$-Si) |

| Figure (panel) | Supplementary Fig. 1d |
|---|---|
| Test used | Kruskal-Wallis test |
| Samples and categories | Current through μLEDs fabricated on LED wafer with FZ-silicon substrate (FZ-Si), μLEDs fabricated on LED wafer with moderately boron-doped silicon substrate ($p^-$-Si), and μLEDs fabricated on LED wafer with heavily boron-doped silicon substrate ($p^+$-Si), at 4 V of forward bias voltage, measured from five different locations on each wafer (B, C, T, L, and R). |
| Statistics provided in figure | Box plots with whiskers and outliers (denoting median, IQR, EVs and outliers) |
| n | 8, 9, 9, 9, & 8 (FZ-Si; B, C, T, L, & R); 9, 9, 9, 8, & 9 ($p^-$-Si; B, C, T, L, & R); & 7, 9, 9, 9, & 9 ($p^+$-Si; B, C, T, L, & R) |
| p-values | $2.67 \times 10^{-2}$ (FZ-Si), $5.51 \times 10^{-2}$ ($p^-$-Si), & $7.90 \times 10^{-1}$ ($p^+$-Si) |
| Other value | $\chi^2 = 1.63 \times 10^1$ (FZ-Si), 9.25 ($p^-$-Si), & 1.70 ($p^+$-Si) |

| Figure (panel) | Supplementary Fig. 1e |
|---|---|
| Test used | Kruskal-Wallis test |
| Samples and categories | Radiant flux generated from μLEDs fabricated on LED wafer with FZ-silicon substrate (FZ-Si), LED wafer with moderately boron-doped silicon substrate ($p^-$-Si), and LED wafer with heavily boron-doped silicon substrate ($p^+$-Si), at 4 V of forward bias voltage, measured from five different locations on each wafer (B, C, T, L, and R). |
| Statistics provided in figure | Box plots with whiskers and outliers (denoting median, IQR, EVs and outliers) |
| n | 8, 9, 9, 9, & 8 (FZ-Si; B, C, T, L, & R); 9, 9, 9, 8, & 9 ($p^-$-Si; B, C, T, L, & R); & 7, 9, 9, 9, & 9 ($p^+$-Si; B, C, T, L, & R) |
| p-values | $6.44 \times 10^{-4}$ (FZ-Si), $1.52 \times 10^{-3}$ ($p^-$-Si), & $3.11 \times 10^{-1}$ ($p^+$-Si) |
| Other values | $\chi^2 = 1.94 \times 10^1$ (FZ-Si), $1.75 \times 10^1$ ($p^-$-Si), & 4.78 ($p^+$-Si) |

| | |
|---|---|
| **Figure (panel)** | **Supplementary Fig. 1f** |
| **Test used** | Kruskal-Wallis test |
| **Samples and categories** | Maximum plug efficiency of μLEDs fabricated on LED wafer with FZ-silicon substrate (FZ-Si), μLEDs fabricated on LED wafer with moderately boron-doped silicon substrate ($p^-$-Si), and μLEDs fabricated on LED wafer with heavily boron-doped silicon substrate ($p^+$-Si), measured from five different locations on each wafer (B, C, T, L, and R). |
| **Statistics provided in figure** | Box plots with whiskers and outliers (denoting median, IQR, EVs and outliers) |
| **n** | 8, 9, 9, 9, & 8 (FZ-Si; B, C, T, L, & R); 9, 9, 9, 8, & 9 ($p^-$-Si; B, C, T, L, & R); & 7, 9, 9, 9, & 9 ($p^+$-Si; B, C, T, L, & R) |
| **p-values** | $7.84 \times 10^{-4}$ (FZ-Si), $3.16 \times 10^{-4}$ ($p^-$-Si), & $1.47 \times 10^{-5}$ ($p^+$-Si) |
| **Other values** | $\chi^2 = 1.90 \times 10^1$ (FZ-Si), $2.10 \times 10^1$ ($p^-$-Si), & $2.76 \times 10^1$ ($p^+$-Si) |

| | |
|---|---|
| **Figure (panel)** | **Supplementary Fig. 3c** |
| **Test used** | Mann-Whitney U test with Bonferroni correction |
| **Samples and categories** | Peak-to-peak magnitudes of PV-induced voltage signal, recorded from all sites on shielded μLED optoelectrodes fabricated using LED wafer with FZ-silicon substrate (FZ-Si), shielded μLED optoelectrodes fabricated using LED wafer with moderately boron-doped silicon substrate ($p^-$-Si), and shielded μLED optoelectrodes fabricated using LED wafer with heavily boron-doped silicon substrate ($p^+$-Si) |
| **Statistics provided in figure** | Box plots with whiskers (denoting median, IQR, and EVs) |
| **n** | 55 (FZ-Si), 49 ($p^-$-Si), & 56 ($p^+$-Si) |
| **p-values** | $1.76 \times 10^{-18}$ (FZ-Si vs. $p^-$-Si), $1.86 \times 10^{-18}$ (FZ-Si vs. $p^+$-Si), & $1.26 \times 10^{-18}$ ($p^-$-Si vs. $p^+$-Si) |
| **Other values** | z = -8.77 (FZ-Si vs. $p^-$-Si), 9.02 (FZ-Si vs. $p^+$-Si), & 8.81 ($p^-$-Si vs. $p^+$-Si) |

| | |
|---|---|
| **Figure (panel)** | **Supplementary Fig. 4c** |
| **Test used** | Mann-Whitney U test with Bonferroni correction |
| **Samples and categories** | Peak-to-peak magnitudes of PV-induced voltage signal, recorded from all sites on shielded μLED optoelectrodes fabricated using LED wafer with heavily boron-doped silicon substrate ($p^+$-Si), electrode arrays fabricated using soda-lime glass substrate (G), and electrode arrays fabricated using LED-on-sapphire substrate (S) |
| **Statistics provided in figure** | Box plots with whiskers (denoting median, IQR, and EVs) |
| **n** | 56 ($p^+$-Si), 20 (G), & 26 (S) |
| **p-values** | $4.11 \times 10^{-11}$ ($p^+$-Si vs. G), $4.19 \times 10^{-13}$ ($p^+$-Si vs. S), & $2.60 \times 10^{-2}$ (G vs. S) |
| **Other values** | z = 6.60 ($p^+$-Si vs. G), 7.25 ($p^+$-Si vs. S), & -2.23 (G vs. S) |

| Figure (panel) | Supplementary Fig. 6b |
|---|---|
| **Test used** | Mann-Whitney U test with Bonferroni correction |
| **Samples and categories** | Peak-to-peak magnitudes of stimulation artifact, recorded from all sites on control μLED optoelectrodes (i.e. non-shielded optoelectrodes with moderate boron doping of the silicon substrate, 1ML), μLED optoelectrodes with shielding layer and moderately boron-doped the silicon substrate (SO), and miniSTAR optoelectrodes. |
| **Statistics provided in figure** | Box plots with whiskers and outliers (denoting median, IQR, EVs and outliers) |
| **n** | 75 (1ML), 67 (SO), & 151 (miniSTAR) |
| **p-values** | $2.71 \times 10^{-23}$ (1ML vs. SO), $2.93 \times 10^{-34}$ (1ML vs. miniSTAR), & $5.64 \times 10^{-24}$ (SO vs. miniSTAR) |
| **Other values** | z = 9.94 (1ML vs. SO), 122.05 (1ML vs. miniSTAR), & 100.98 (SO vs. miniSTAR) |

| Figure (panel) | Supplementary Fig. 9c |
|---|---|
| **Test used** | Kruskal-Wallis test |
| **Samples and categories** | Peak-to-peak magnitudes of stimulation artifact, recorded from two sites at the bottom of each shank (sites 1 & 2) on miniSTAR optoelectrodes during LED driving with current pulses different shapes – trapezoidal, sinusoidal, and sigmoidal – with 10 - 90 % rise times of approximately 1 ms. |
| **Statistics provided in figure** | Box plots with whiskers (denoting median, IQR, and EVs) |
| **n** | 18 (for all categories) |
| **p-value** | $9.33 \times 10^{-2}$ |
| **Other values** | $\chi^2 = 4.74$ |

### Supplementary Figures

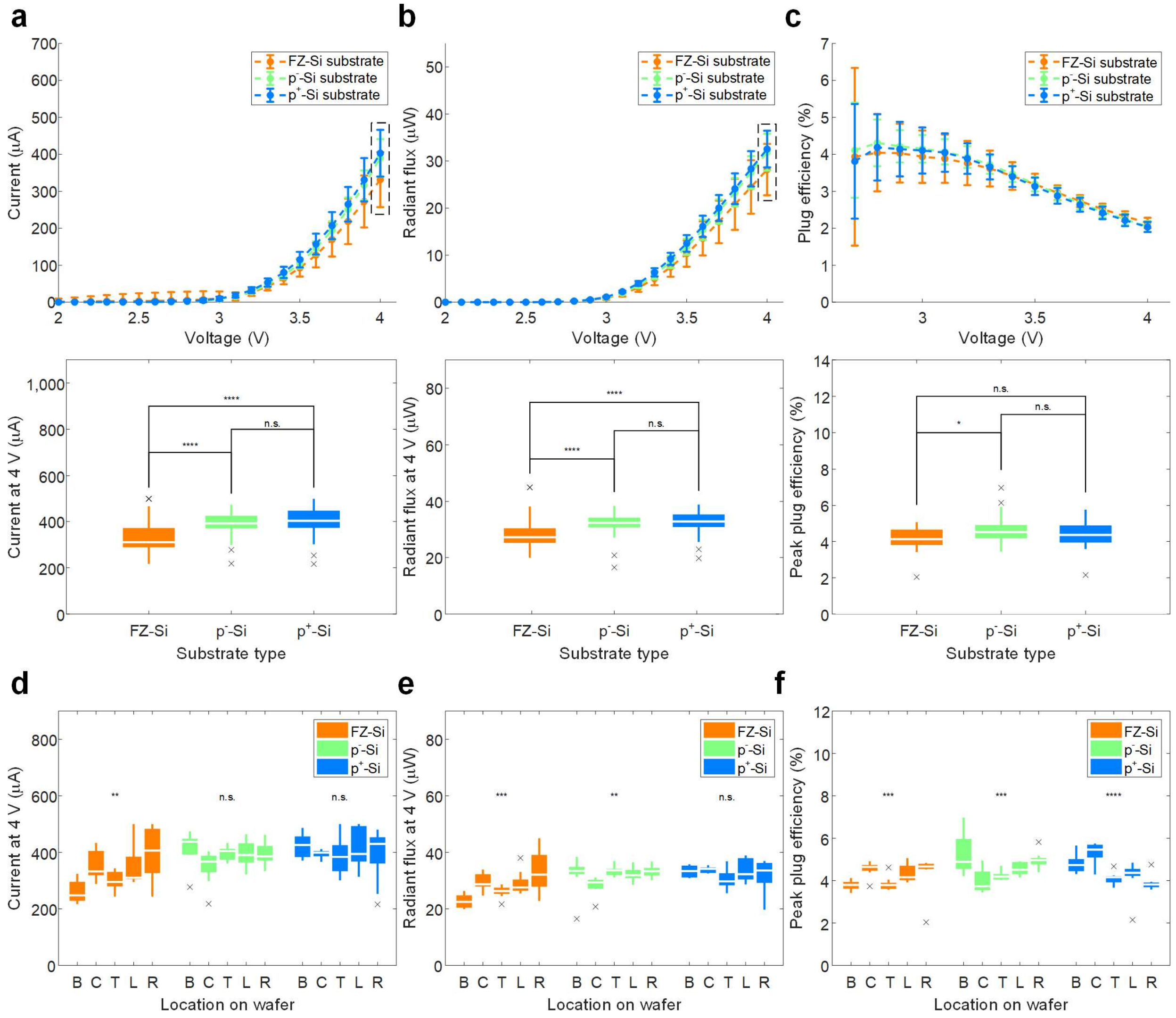


**Supplementary Figure 1: Electrical and optical characteristics of μLEDs fabricated using GaN-on-Si LED wafers with differently boron doped silicon substrates.**

(**a**) Mean μLED forward current at different bias voltages. Distribution of current from LEDs fabricated using wafers with different substrate doping densities at 4 V of forward bias voltage are shown at the bottom. At the top, error bars indicate one standard deviation. At the bottom, boxes indicate interquartile ranges, white lines medians, whiskers non-outlier extreme values and x marks outliers. (**b**) Mean μLED output radiant flux at different bias voltages and their distribution at 4 V of forward bias voltage. (**c**) Mean μLED plug efficiency at different bias voltages and the distribution of the peak plug efficiency. (**d**) Distribution of the current from LEDs fabricated on different locations on the wafer at 4 V of forward bias voltage. (**e**) Distribution of the radiant flux from LEDs fabricated on different locations on the wafer at 4 V of forward bias voltage. (f) Distribution of the peak plug efficiency of LEDs fabricated on different locations on the wafer. Results of statistical tests are provided in Supplementary Table 2.

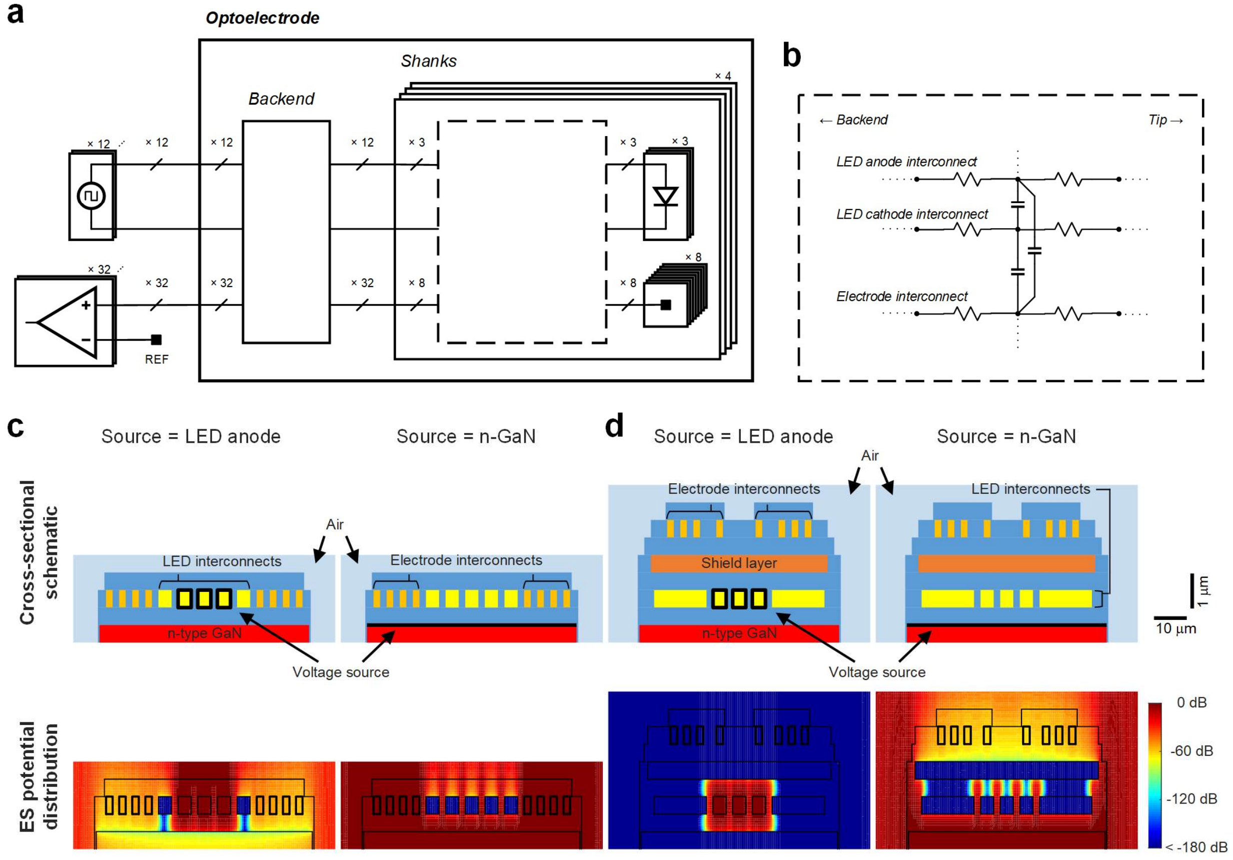


**Supplementary Figure 2: Generation of EMI-induced stimulation artifact.** (**a**) System-level electrical circuit schematic diagram of μLED optoelectrode, LED driving system, and neural signal recording system. Some details, including some resistors representing the line and the contact resistances, are omitted for better visibility. The equivalent circuit of the backend, which is similar to that of the shank shown in part b, is also omitted. (**b**) Simplified electrical circuit schematic diagram of a shank of μLED optoelectrode. Only one of each type of interconnect is shown for better visibility, and inductors were ignored due to their small values. (**c**) Results of finite-element-method (FEM) simulation of electrostatic potential distribution inside the one-metal-layer (non-shielded) μLED optoelectrode shank cross-section due to voltage from different EMI sources. Regions in dark light blue inside the air indicate silicon dioxide insulators. Sources of EMI are highlighted with bold black lines. Electrostatic potentials of the highlighted regions were set as 1 V, while those of the other parts of the LED driving circuitry were set as

0 V. (**d**) Results of FEM simulation of electrostatic potential distribution inside shielded optoelectrode shank cross-section due to voltage from different EMI sources.

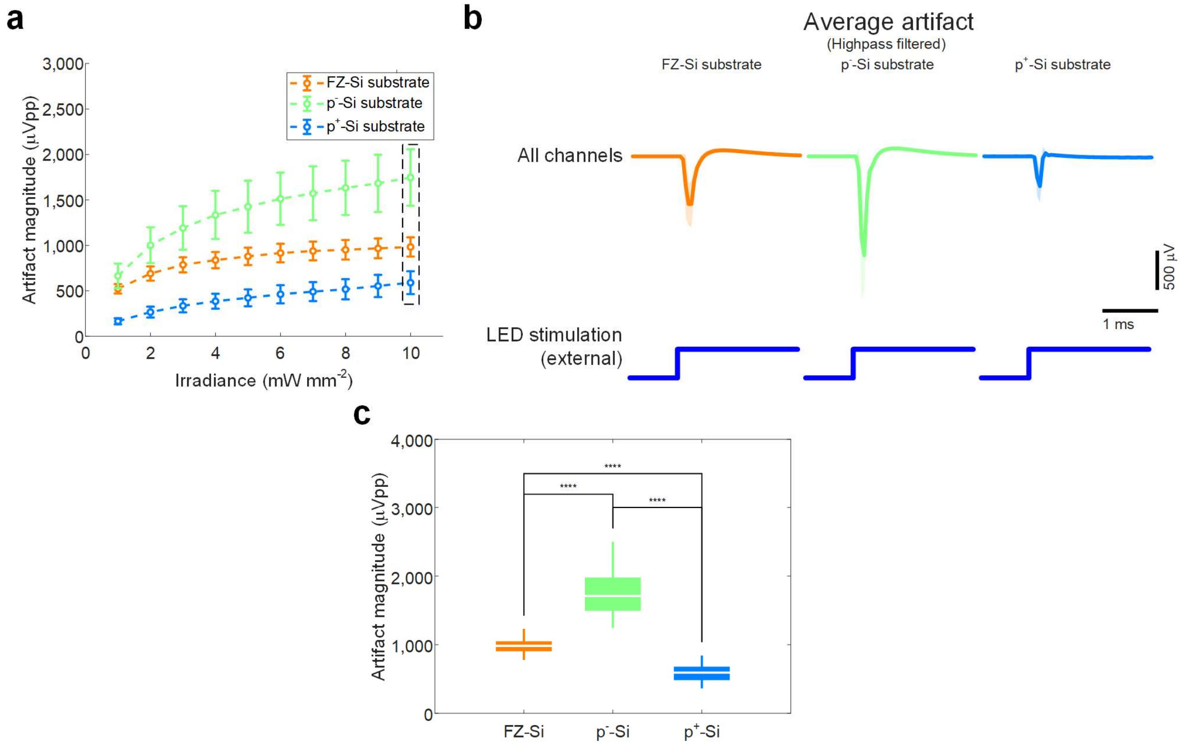


**Supplementary Figure 3: PV-induced voltage signals measured on shielded μLED optoelectrodes fabricated using GaN-on-Si LED wafers with differently boron-doped silicon substrates.** (a) Comparison of mean peak-to-peak magnitude of highpass filtered voltage signal recorded on shielded μLED optoelectrodes with different substrate doping densities upon external light exposure. Error bars indicate one standard deviation. (**b**) Mean highpass filtered waveforms whose mean peak-to-peak magnitudes are shown in part **a**, inside the rectangle with black dashed sides. Shaded regions show one standard deviation away from mean. (**c**) Peak-to-peak magnitudes of highpass filtered voltage signal whose mean waveforms are shown in part **b**. Boxes indicate interquartile ranges, white lines medians, and whiskers extreme values. Mean (± SD) peak-to-peak magnitudes are 982.43 (± 105.76), 1746.80 (± 310.89), and 589.72 (± 125.64) μVpp for devices with FZ-Si substrate (n = 55), $p^-$-Si substrate (n = 49), and $p^+$-Si substrate (n = 56), respectively. Results of statistical tests are provided in Supplementary Table 2.

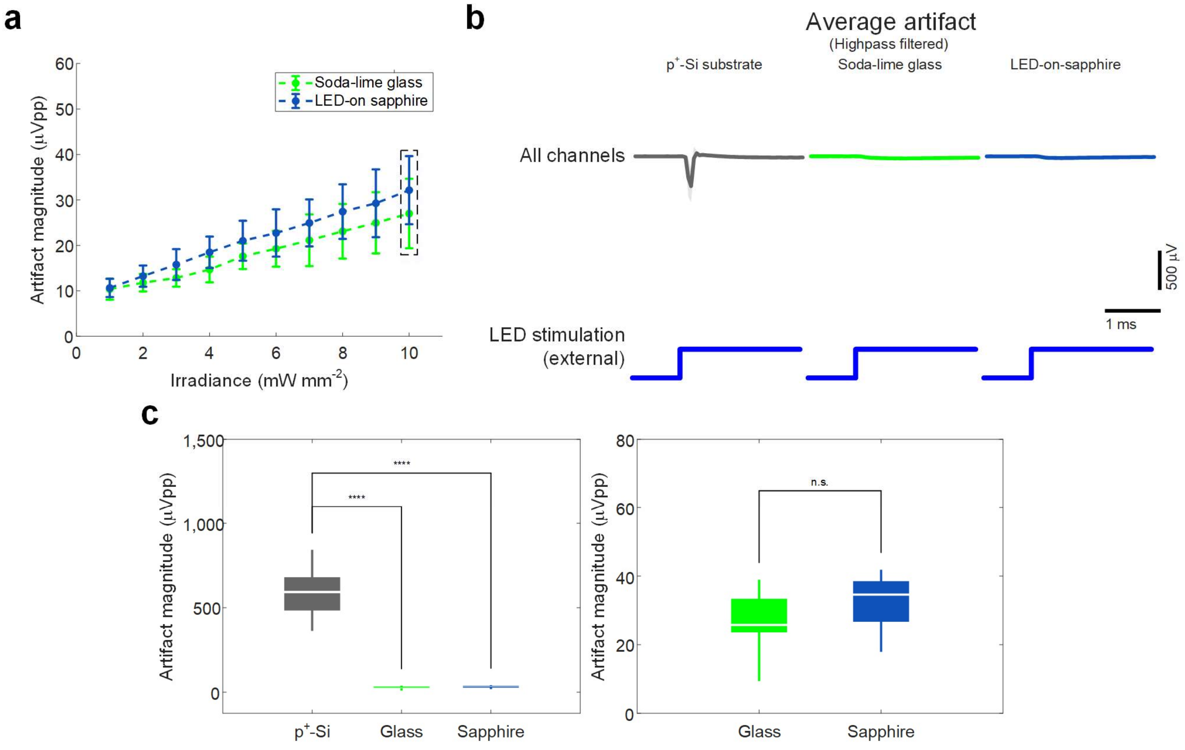


**Supplementary Figure 4: PV-induced voltage signals measured on electrode arrays fabricated using non-silicon substrates.** (a) Comparison of mean peak-to-peak magnitude of highpass filtered voltage signal recorded on electrode arrays fabricated with different substrates upon light exposure. Error bars indicate one standard deviation. (**b**) Mean highpass filtered waveforms whose mean peak-to-peak magnitudes are shown in part **a**, inside the rectangle with black dashed sides. Mean highpass filtered waveform of voltage signal recorded from shielded μLED optoelectrodes with heavily boron doped-silicon substrate is shown for comparison. Shaded regions show one standard deviation away from mean. (**c**) Peak-to-peak magnitudes of the highpass filtered voltage signal whose mean waveforms are shown in part **b**. Boxes indicate interquartile ranges, white lines medians, and whiskers extreme values. Mean (± SD) peak-to-peak magnitudes are 589.72 (± 125.64), 27.00 (± 7.64), and 32.14 (± 7.49) μVpp for optoelectrodes with p$^+$-Si substrate (n = 56), electrode arrays fabricated using soda-lime glass substrate (n = 20), and electrode arrays fabricated using LED-on-sapphire substrate (n = 26), respectively. Results of statistical tests are provided in Supplementary Table 2.

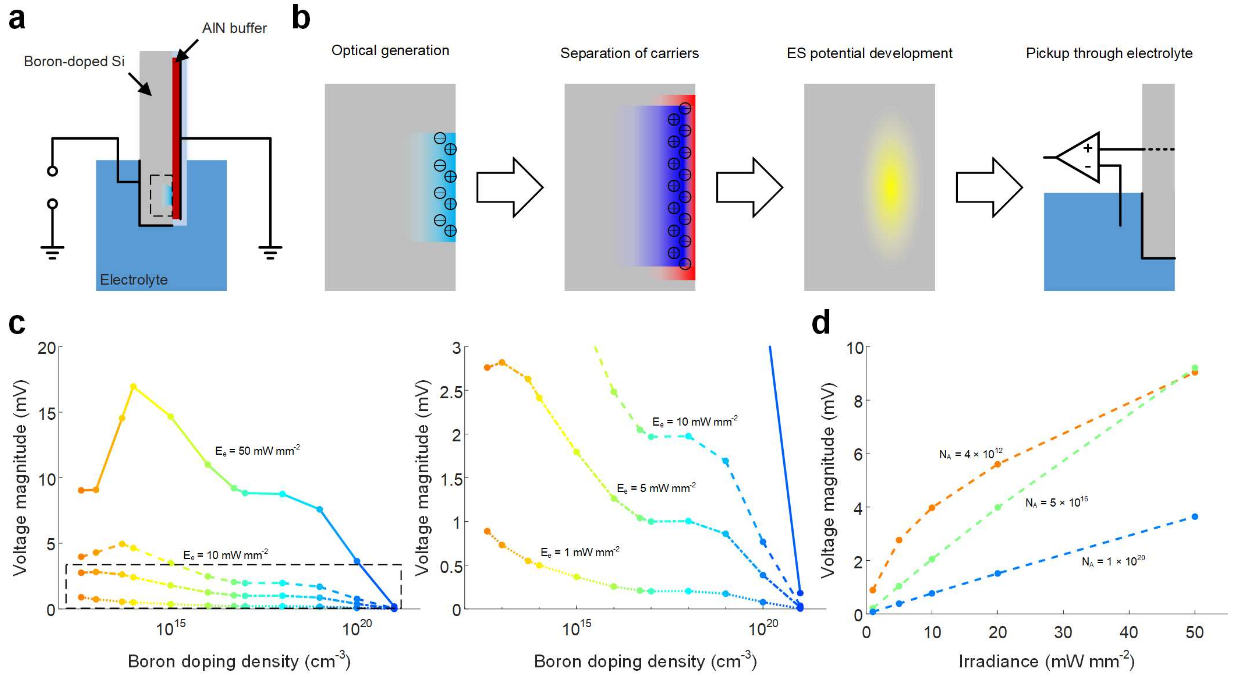


**Supplementary Figure 5: Generation of PV-induced electrostatic potential and consequent stimulation artifact.** (**a**) Schematic illustration of the cross-section of the 3D model used in finite-element-method (FEM) device physics simulation. Light blue region indicates silicon dioxide insulator. The electrolyte was taken into account by applying a boundary condition to the electrode at the interface. Bold black lines indicate electrodes in the model and their boundary conditions. (**b**) Schematic illustrations of the processes through which electrostatic potential is induced and PV-induced stimulation artifact is generated. The first three panels are the magnified views of the region inside the rectangle with the black dashed sides on part **a**. Circles with plus signs indicate holes, circles with minus signs electrons, shading in light blue optical generation, shading in blue distribution of holes, shading in red distribution of electrons, and shading in yellow electrostatic potential. In steady state, all the processes occur simultaneously and in turn maintains a steady distribution of electrostatic potential inside the substrate. (**c**) Steady-state voltage of the substrate-electrolyte interface, calculated at different doping densities and light intensities. The right panel shows the magnified view of the region inside the rectangle with black dashed sides on the left panel. (**d**) Steady-state substrate-electrolyte interface voltage of substrates with a few selected boron doping densities at different light intensities.

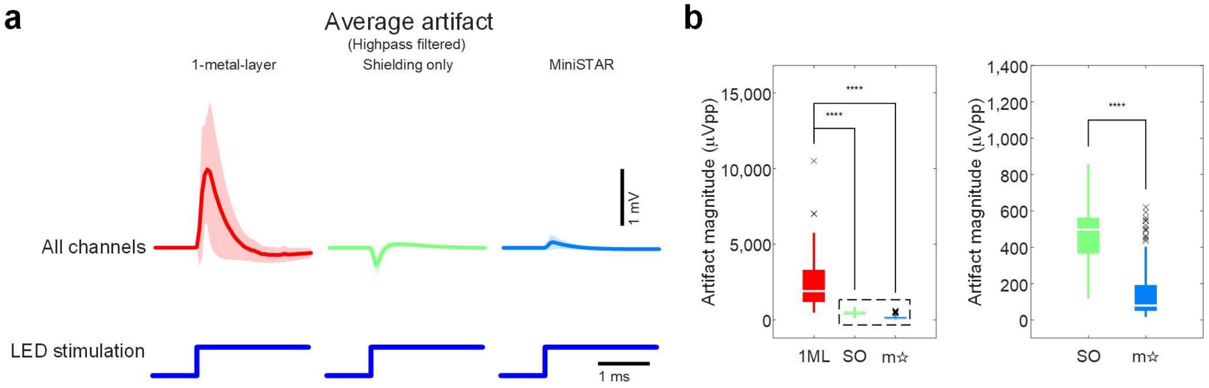


**Supplementary Figure 6: Reduction of stimulation artifact in miniSTAR μLED optoelectrodes.** (**a**) Mean highpass filtered waveforms of stimulation artifact recorded from different μLED optoelectrodes. Each mean was calculated using only the signals recorded from channels that correspond to electrodes on the shank on which a μLED was turned on. LED drive signal with resulting LED surface irradiance of 75 mW mm$^{-2}$ was used. Shaded regions show one standard deviation away from mean. (**b**) Peak-to-peak magnitudes of the signals whose averages are plotted in part **a**. The right panel shows the magnified view of the region inside the rectangle with black dashed sides on the left panel. Boxes indicate interquartile ranges, white lines medians, whiskers non-outlier extreme values, and black x marks outliers. Mean (± SD) peak-to-peak magnitudes are 2477.75 (± 1733.83), 474.59 (± 146.26), and 146.05 (± 143.40) μVpp for one-metal-layer devices (n = 75), shielded devices with no substrate doping modification (n = 67), and miniSTAR devices (n = 151), respectively. Results of statistical tests are provided in Supplementary Table 2.

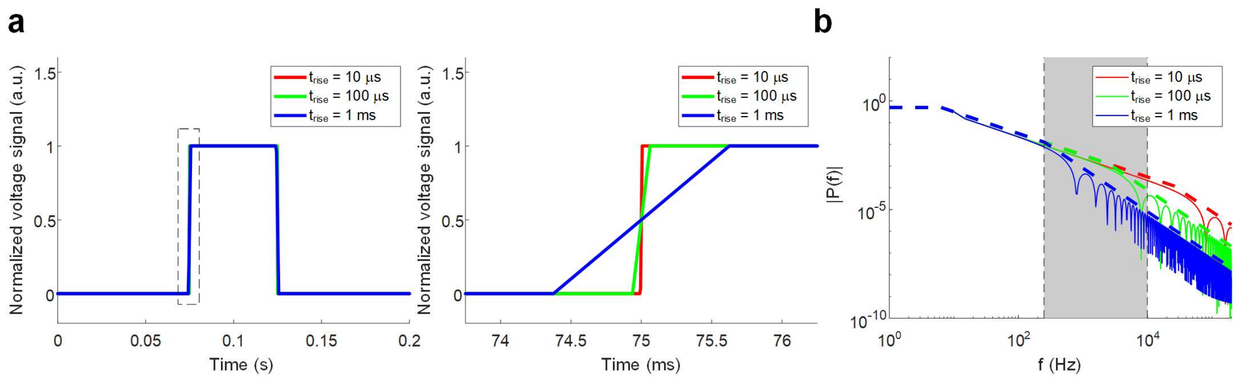


**Supplementary Figure 7: Voltage pulses with different rise times and their spectra.** (**a**) Time-domain plot of voltage pulses with rise times (10 - 90 % rise times)of 10 μs, 100 μs, and 1 ms. The right panel shows the magnified view of the region inside the rectangle with black dashed sides on the left panel. (**b**) Frequency spectrum of pulses shown in part **a**, showing both envelopes (in dashed lines) and values evaluated at a few selected harmonic frequencies (in thin solid lines. Only prime numbered harmonics (f = 5 × (2, 3, 5, 7, etc.)) are shown for better visibility). Frequency range of 250 Hz < f < 10 kHz is highlight with a shade of grey.

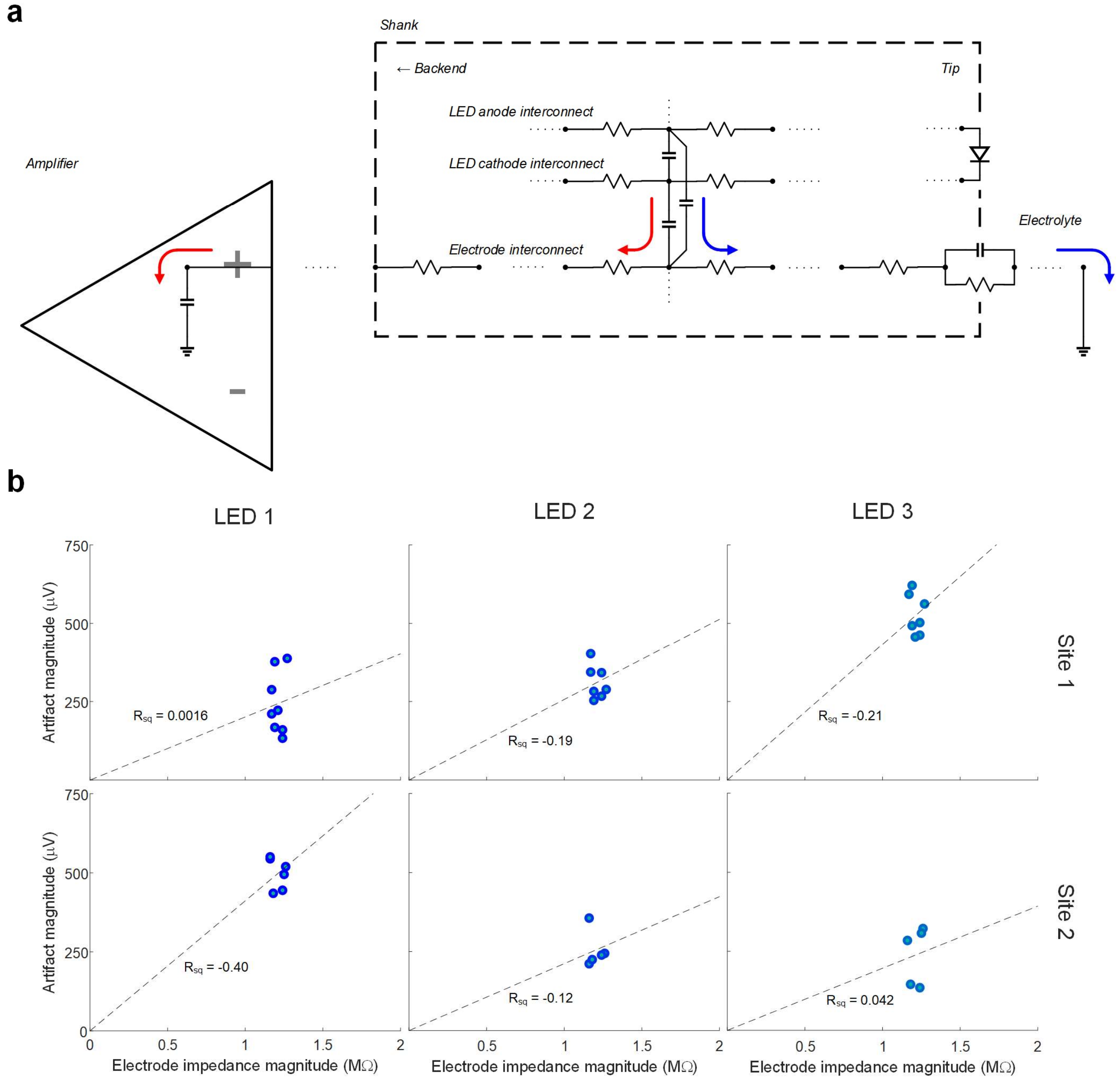


**Supplementary Figure 8: Relationship between the recording electrode impedance and the stimulation artifact magnitude.** (**a**) Detailed equivalent circuit diagram of the neural signal recording circuitry, whose simplified version is presented in Supplementary Figure 2b. A parallel RC component, representing the electrical double layer at the electrode / electrolyte interface, and a capacitor, representing the signal amplifier's input impedance, are shown at the ends of the series resistor components representing the recording electrode interconnect. Division of the capacitively coupled current between the two branches is visualized with two arrows with different colors. **(b)** Peak-to-peak magnitudes of the highpass filtered stimulation artifact resulting from rectangular LED input voltage signals with 3.5 V peak-to-peak amplitude ($V_{low\text{-}level}$ = 0 V, $t_{rise}$ = 5 ns), recorded from the sites 1 and 2 (the bottommost sites) on miniSTAR μLED optoelectrodes. All impedances are evaluated at 1 kHz. Small $R_{sq}$ values suggest lack of clear correlation between the artifact magnitude and the electrode impedance.

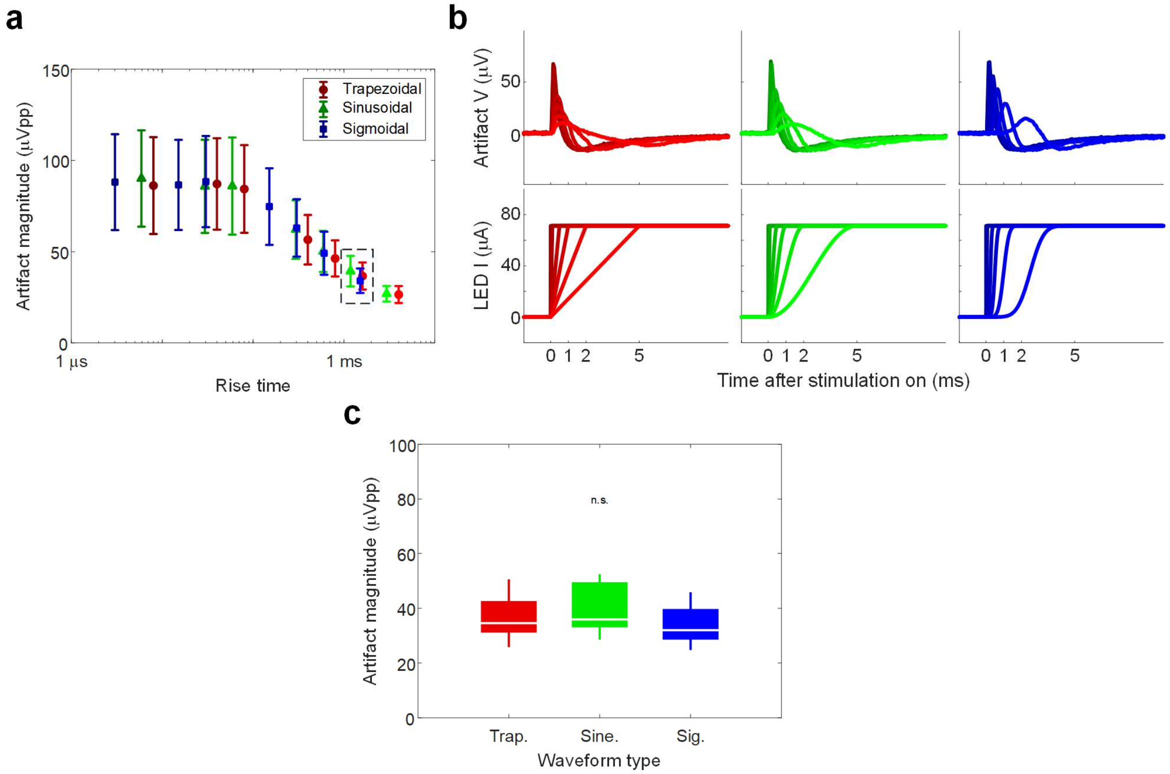


**Supplementary Figure 9: Artifact resulting from stimulation with current-driven LEDs.**

(**b**) Mean peak-to-peak magnitude of highpass filtered stimulation artifact recorded on miniSTAR μLED optoelectrodes from channels indicated in Fig 5**a**. X coordinates indicate the 10 - 90 % rise time of the pulse. Error bars indicate one standard deviation. (**b**) Mean waveforms of recorded stimulation artifact, whose mean peak-to-peak magnitudes are shown in part **a**, and their input current signals. Stimulation artifact resulting from an input current signal is indicated with the same color. **(d)** Peak-to-peak magnitudes of highpass filtered stimulation artifact for a few selected conditions whose means are shown inside the rectangle with black dashed sides in part **a**. Boxes indicate interquartile ranges, white lines medians, and whiskers extreme values. Mean (± SD) peak-to-peak magnitudes are 36.68 (± 7.41), 39.32 (± 8.36), and 34.09 (± 6.71) μVpp, from left to right. Results of statistical tests are provided in Supplementary Table 2.
